# Efficacy of a long lasting ivermectin formulation against malaria vectors in a semi-field trial in Bapla, Southwest of Burkina Faso

**DOI:** 10.64898/2026.07.31.741508

**Authors:** Cheick Oumar W. Ouédraogo, Fabrice A. Somé, Samuel Beneteau, Emmanuel Sougué, Angélique Porciani, Dieudonné D. Soma, Nicolas Moiroux, Sié Rodrigue Dah, Sié Hermann Pooda, André B. Sagna, Cédric Pennetier, Lamidi Zela, Mady Ndiaye, Sophie Le Lamer-Déchamps, El Hadji A. Niang, Karine Mouline, Roch K. Dabiré

## Abstract

**Background:** Ivermectin, a widely used endectocide with activity against *Anopheles (An)* mosquitoes, has emerged as a promising complementary tool for malaria vector control. However, currently approved oral formulations provide only short-lived plasma concentrations at mosquitocidal levels, thereby limiting their operational impact. To overcome this limitation, a long-acting injectable formulation based on ivermectin and BEPO technology (mdc-STM-001) was selected through laboratory screening using mosquito colonies. This formulation demonstrated mosquitocidal efficacy meeting WHO criteria, with a hazard ratio (HR) >4 maintained for at least two months. The present study aims to further characterize the efficacy of ivermectin under robust field conditions and to expand susceptibility assessments to wild *Anopheles* populations representing different species and vectorial status. Ivermectin was tested in cattle at two doses under field conditions using experimental huts and tent traps. Plasma drug levels were also monitored to characterize the pharmacokinetic profile of ivermectin.

**Methods:** Thirty-six calves were involved in the study and allocated to three groups (n = 12 per group). The mdc-STM-1.0 group received a single subcutaneous injection of the long-acting ivermectin formulation mdc-STM-001 at a dose of 1.0 mg/kg; the mdc-STM-1.5 group received the same formulation at 1.5 mg/kg and the third group served as controls. Using a three-arm design, cattle from each treatment group were simultaneously exposed to mosquitoes in experimental huts and tent traps, following a Greco-Latin Square rotation to minimize positional confounding factors. Pharmacokinetic data were collected to assess ivermectin release from the subcutaneous depot. Correlations between plasma ivermectin concentrations and mosquito mortality were explored, and survival analysis including Kaplan-Meier and cox proportional hazards models were performed to assess the effect of treatment on mosquito survival.

**Results:** The mdc-STM-001 formulation was well tolerated and produced sustained, dose-dependent plasma ivermectin concentrations, with detectable levels maintained for more than 90 days after a single administration. Both doses induced strong mosquitocidal activity against wild *Anopheles,* with hazard ratios reaching 8.03 and 16.47 for the 1.0 and 1.5 mg/kg doses, respectively. *An. funestus* and *An. rufipes* were the most susceptible species (LC50: 5.1-6.9 ng/mL) exhibiting prolonged LC50 coverage (>75 days), whereas *An. coustani* was less susceptible (LC50: 12.4-19.2 ng/mL) resulting in a shorter duration of effective coverage (17-41 days).

**Conclusions:** Taken together, our results revealed the potential of the mdc-STM-001 long-acting ivermectin formulation as an effective endectocide against residual malaria transmission by primary and secondary vectors and establish evidence basis to support its progression to early-phase clinical trials in humans.

## Background

Vector-borne diseases (VBD) remain a major global public health challenge accounting for more than 17% of all communicable diseases, with around 80% of the human population at risk of infection [1]. Among all VBD, malaria is the most prevalent and debilitating parasitic disease, causing approximately in 2024, 282 million clinical cases worldwide with 610,000 deaths, the majority occurring in Sub-Saharan Africa [2]. In Burkina Faso, despite sustained control efforts, the country is still among the most affected, ranking 8^th^ globally with 8,324,000 malaria cases, and 16,184 deaths [2]. One may assume that the multi-resistance profile of malaria vectors to the common insecticides can constitute the major threat to the effectiveness of vectors control interventions [3–7]. Current vector control strategies, including Insecticide Treated Nets (ITNs) and Indoor Residual Spraying (IRS), primarily target *Anopheles* mosquitoes that feed and rest indoors. While these tools have reduced malaria transmission, their effectiveness is limited against mosquito populations exhibiting exophagic and exophilic behaviors. And then some *Anopheles* mosquitoes are naturally more zoophilic and readily exploit livestock as alternative blood meal sources [8]. Such feeding plasticity can contribute to the persistence of mosquito populations, particularly in areas where human hosts are less accessible, being protected by ITNs [9, 10]. In addition, sustained exposure to indoor-based interventions may select behavioral adaptations that reduce mosquito contact with humans indoors, promoting increased outdoor host-seeking and feeding activity. Consequently, strategies targeting only mosquito feeding and resting indoors may fail to affect a substantial fraction of the vector population capable of sustaining malaria transmission.

To address these gaps, innovative complementary vector control tools are needed to target mosquito populations that continue to sustain malaria transmission despite the widespread use of conventional interventions. In this context, ivermectin (IVM), a well-known antiparasitic drug, has recently emerged as a promising complementary tool for vector control.

Traditionally administered to treat parasitic infections in humans and livestock since decades [11, 12], ivermectin has demonstrated recently a broad range of biological activities, including antiviral, anticancer, anthelmintic effects at different concentrations [13]. Importantly, accumulating evidence indicates that ivermectin also exerts potent mosquitocidal activities by reducing mosquito survival after blood feeding on treated hosts [8, 14]. Beyond its direct lethal effects, ivermectin induces some sublethal effects in surviving mosquitoes such as reduced fecundity and fertility[15–17]. These findings have therefore prompted the evaluation of ivermectin as a complementary malaria vector control intervention through mass drug administration (MDA) strategy. Some cluster-randomized trials demonstrated that ivermectin MDA can reduce mosquito survival and malaria transmission in endemic settings. In Burkina Faso, the RIMDAMAL trial showed that repeated ivermectin administration reduce the incidence of malaria episodes in children [18]. More recently, BOHEMIA trial in Kenya confirmed that ivermectin administered alongside standard malaria control interventions can reduce malaria incidence by 26% in children’s populations aged 5-15 years old [19]. An important limitation identified in these trials was the shorter than expected duration of ivermectin’s mosquitocidal effect, even at higher doses [20]. Consequently, multiple rounds of MDA would be required to maintain effective mosquitocidal plasma concentrations over time [21–23]. To overcome this limitation, long-acting injectable ivermectin formulations (LAIF) based on the BEPO**^®^** technology have been developed to provide sustained drug release over several weeks to months [24, 25]. By maintaining insecticidal plasma concentrations for extended periods, these formulations have the potential to substantially enhance the impact of ivermectin-based strategies for malaria control. Preclinical evaluation of three LAIF candidates in a cattle model identified the mdc STM 001 formulation as the most promising candidate for clinical development. A single subcutaneous injection at 0.6 mg/kg maintained mosquitocidal drug concentrations, achieving hazard ratios above 4 against both susceptible and resistant Anopheles populations for at least 2 months, thereby meeting WHO Preferred Product Characteristics (PPC) for malaria endectocides.

In the present study, we evaluated the pharmacokinetics, pharmacodynamics (PK/PD) and safety of two higher doses of mdc-STM-001 (1.0 and 1.5 mg/kg). The mosquitocidal activity of both doses was assessed under field conditions using wild *Anopheles* populations from Bapla, southwestern Burkina Faso. This study also provided the first opportunity to evaluate the efficacy of long-acting ivermectin against both primary and secondary malaria vector species, potentially contributing to residual *Plasmodium* transmission. The findings generate essential preclinical evidence to support progression to a first-in-human Phase 1 clinical trial.

## Materials and methods

### Study site

The field trials were carried out between September 2021 and January 2022 in Bapla village (10°53’39.34”N; 3°15’46.68”O), located in the Sud-Ouest Region of Burkina Faso, approximately 10 km from Diébougou. The climate is Sudano-Sahelian, characterized by two distinct seasons: a dry period from November to May, followed by a rainy season from May to November with the mean annual precipitation averaging 1051 mm. The primary economic activities are agriculture and fishing, favorized by the presence of an hydro-agricultural dam [26]. Malaria transmission is primarily driven by *Anopheles gambiae*, *Anopheles coluzzii*, *Anopheles funestus s.s*, and *Anopheles arabiensis*, which exhibit marked seasonal dynamics [27]. The village is characterized by wooded savannah vegetation surrounded by diverse natural habitats, including gallery forests and both temporary and permanent water bodies, creating favorable conditions for mosquitoes’ proliferation.

### Overall entomological study design

Two complementary approaches of calve exposure to wild *Anopheles* mosquitoes were used to be able to apprehend the great diversity of species present in the area, with respect to their biting proclivities, using either i) experimental hut (or “cases-pièges”), recognized by the WHO as a phase II insecticide evaluation tool [28], to target indoor biting mosquitoes and also to characterize the potential deterrent or attractant effect of treated calves, or ii) “tent trap” (*i.e.,* untreated mosquito net) to capture outdoor biting and/or zoophagic mosquitoes. In the study area, malaria vectors exhibit marked trophic plasticity. A high proportion of *Anopheles gambiae*, *Anopheles coluzzii*, and *Anopheles arabiensis* are known to feed not only on humans but also on domestic animals, with a pronounced preference for cattle [29, 30].

Therefore, despite using cattle as bait, we were confident that we would capture the major vector species driving malaria transmission in the area and measure key, representative entomological outcomes related to ivermectin susceptibility for these species.

The combined use of experimental huts and tent traps enabled the investigation of both endophagic and exophagic vector populations, respectively. A graphical overview of the study protocol is provided in **Fig. 1**. Two LAIF doses were tested: 1.0 and 1.5 mg/kg single dosing performed at different days in each round for easier logistics. Thirty-six calves in total were involved in this study, 18 for each of the approach, forming 2 study groups (hut and tent). In each of the study groups, three treatment groups were randomly constituted using stratified randomization to balance body weight across groups: 6 cattle received mdc-STM-001 at the dose of 1.0 mg/kg, 6 cattle at 1.5 mg/kg, and 6 cattle were left untreated as controls. The treatments were administered in a staggered manner at one-week intervals to facilitate rotations between the huts or tents (n=6 for each approach). Each rotation group comprised two cattle per treatment, all treatments were administered simultaneously. Accordingly, three rotation groups were established: Group R1 was treated at t0, Group R2 seven days later, and Group R3 fifteen days after Group R1. Cattle from each rotation group was exposed nightly to wild mosquitoes, with each cattle rotating between huts for 6 nights following a “carré gréco-latin” design, allowing each calf to spend a night in each hut or tent. Rotations were performed between 2-8 (round 1), 30-36 (round 2), 60-66 (round 3) and 90-96 (round 4) days after treatment. Each round represented therefore 12 cattle-nights per treatment.

**Fig. 1.**
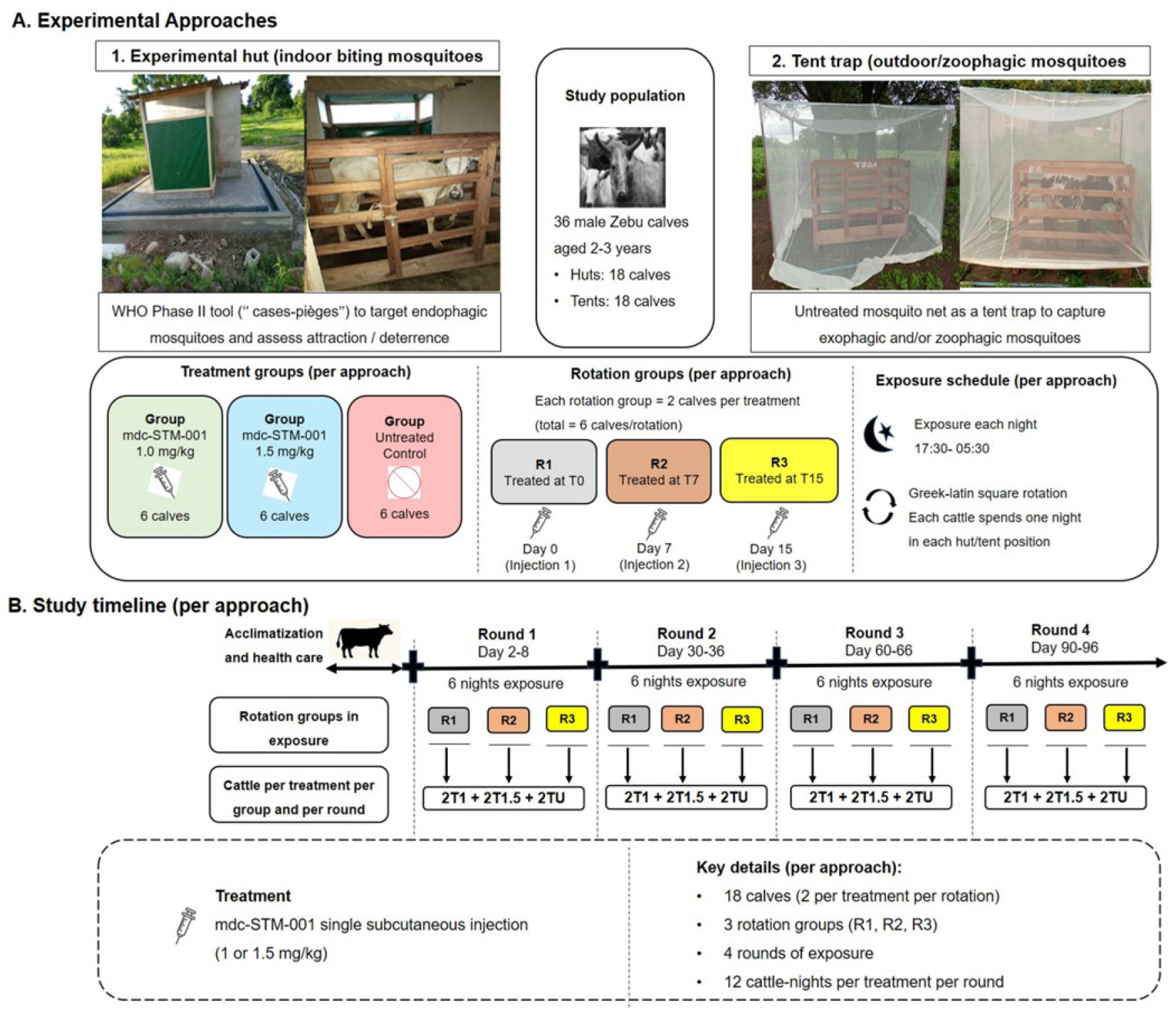
Experimental overview of means and schedule of cattle exposure to mosquitoes. A. Pictures of the two experimental devices used for cattle exposure: experimental hut and tent trap. The layout for constituting treatment and rotation groups is given for both approaches below the pictures. B. Schedule for pre-treatment cattle management, treatment and post-treatment exposure to mosquitoes for each rotation group and each month (round) after injection. T1 = Treated at 1.0 mg/kg of ivermectin, T1.5 = Treated at 1.5 mg/kg of ivermectin, TU = Untreated

### Experimental hut description

Experimental huts are semi-field systems widely used to evaluate vector control tools against wild mosquitoes under conditions that closely mimic natural settings. At Bapla, 6 experimental huts were built near the dam following the typical West african style [31].

In our test system, ivermectin-treated cattle were used as bait instead of habitually human sleepers. Nevertheless, mosquito collection was performed by human collectors following standard procedures. Because the huts were newly constructed, preliminary blank trials were conducted to validate their suitability for mosquito sampling, confirming their effectiveness as trapping systems for evaluating vector control strategies (Unpublished data).

### Tent traps description

Net tent traps are mesh netting enclosures used to capture mosquitoes attracted to animal or human bait under natural field conditions, allowing the study of behaviors such as host preference and outdoor biting [32].

In this study, tents made of untreated mosquito nets (2.80 m x 1.80 m x 1.90 m) were positioned in a fixed configuration outdoors in villagers’ concessions, with an average inter-tent distance of approximately 1000 meters. Cattle were placed inside as bait, and the nets were raised approximately 10cm above the ground to allow mosquito entry.

### Long acting ivermectin formulation using BEPO^®^ technology

The long-acting ivermectin formulation (LAIF) used for cattle treatment is named mdc-STM-001. This formulation has been developed using the BEPO^®^ technology, a proprietary long-acting injectable technology (Medincell, Jacou, France) [33], and was selected as the optimal formulation to be injected to human against malaria vectors in western Africa based on preclinical evaluation [34]. Upon subcutaneous injection, BEPO^®^ forms an in situ bioresorbable polymeric depot *via* a solvent-exchange mechanism, leading to precipitation of a polymer matrix that enables sustained drug release. Each formulation consisted of three components: i) a biocompatible solvent to ensure injectability, ii) ivermectin, as the active pharmaceutical ingredient, and iii) a mixture of two biocompatible and biodegradable polymers that control drug release kinetics. These polymers undergo progressive hydrolysis into bioabsorbable by-products over time, thanks to a simple hydrolysis mechanism.

The mdc-STM-001 LAI formulation was imported into Burkina Faso with clearance from the Direction Générale des Services Vétérinaires of Burkina Faso (Permit No 2020/199) and stored at room temperature, protected from light, until administration to cattle.

### Cattle hosts, formulation injections and coproscopic examination

Thirty-six male zebu calves, aged 2-3 years old with an initial average weight (±SD) of 101.4±13.1 kg were purchased from markets in Bapla and Diebougou surrounding villages, one month prior to the study to allow acclimation. Upon arrival at the enclosure, calves were treated at baseline by a trained veterinarian for trypanosomiasis and gastrointestinal parasites using Berenil 2000^®^ and Benzal^®^, respectively. Animals were fed rice straw and cottonseed cake, with water and sea salt, provided ad libitum. The mdc-STM-001 formulation was administered as a single subcutaneous injection into the loose skin anterior to the shoulder at two doses: 1.0 and 1.5 mg/kg. The 1.5 mg/kg dose represents the upper limit of feasible subcutaneous volumes for human application and was included to assess the dose-dependent effects on pharmacokinetics and mosquitocidal efficacy between

1.0 and 1.5 mg/kg. Nominal injection volumes per cattle group and per approach are provided in Table 1. The animals’ identification numbers, their weight upon arrival at the Bapla pen station and on the day-2 before injection, the administered mdc-STM-001 formulation volume per animal (ranged from 1.05 to 2.28 mL) are detailed in the **supplementary file S1: Table 1**. Feces from experimental calves were collected for coproscopic evaluation of gastrointestinal parasites. Fecal samples (∼ 200 g) were collected directly from rectum of each calf at multiple timepoints (Day 0, 2, 30, 60 and 90) for coproscopic examination. Endoparasite prevalence and infection intensity were assessed using the McMaster and sedimentation methods. *Strongles, Strongyloides* and *Ascaris* eggs were quantified using McMaster technique [35]. Briefly, 3g of feces were diluted in 42 mL of 40% NaCl solution, homogenized, filtered (200 µm) and loaded into a McMaster chamber. Eggs were counted under 10x magnification and expressed as eggs per gram of feces. *Fasciola* and Paramphistom eggs were checked using the sedimentation method after dilution of fecal samples in water (1:15, w/v), filtration and 8 h sedimentation, followed by microscopic examination at 40x magnification. Overall animal health and well-being were also monitored throughout the study by assessing clinical signs, local injection site tolerance, body weight and weight gain.

**Table 1.**
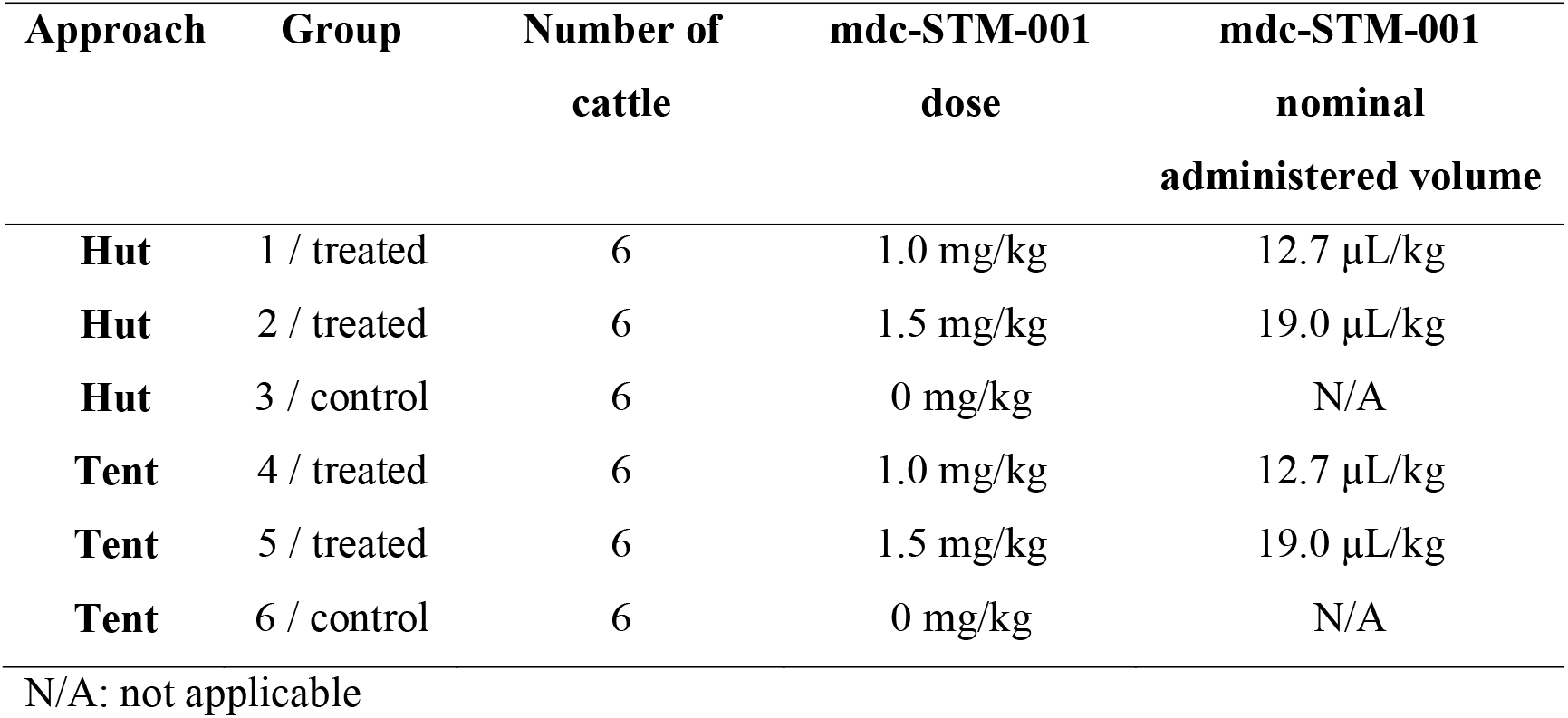
Nominal treatments volumes of mdc-STM-001 ivermectin per group.

| Approach | Group | Number of cattle | mdc-STM-001 dose | mdc-STM-001 nominal administered volume |
| --- | --- | --- | --- | --- |
| Hut | 1 / treated | 6 | 1.0 mg/kg | 12.7 µL/kg |
| Hut | 2 / treated | 6 | 1.5 mg/kg | 19.0 µL/kg |
| Hut | 3 / control | 6 | 0 mg/kg | N/A |
| Tent | 4 / treated | 6 | 1.0 mg/kg | 12.7 µL/kg |
| Tent | 5 / treated | 6 | 1.5 mg/kg | 19.0 µL/kg |
| Tent | 6 / control | 6 | 0 mg/kg | N/A |

N/A: not applicable

In both hut and tent approaches, cattle were placed inside a wooden box during exposure to mosquitoes to limit their movements.

### Procedure for attractiveness proxy measurements and mosquitocidal efficacy assessments of mdc-STM-001 treatments

#### Cattle exposure to mosquitoes

Each rotation group consisted of six cattle including 2 animals per treatment group (**Fig. 1**). Three independent rotation groups were established for each experimental approach (tent and hut). Consequently, each treatment group included six cattle per experimental approach, corresponding to a total of 36 cattle across all treatment groups and experimental approaches combined. Cattle were exposed to wild mosquito populations from 5 :30 pm to 5 :30 am for 6 consecutive nights per month (round) throughout the study period, for a total of four exposure rounds (*i.e.,* 24 exposure nights per animal) (**Fig. 1**). To account for potential positional effects, cattle were rotated from one night to the other so the 6 cattle were exposed to mosquitoes at each tent or hut position. Mosquitoes that blood-fed on exposed cattle were collected as early as possible in the morning (between 5:30 and 7:00 am), prior to temperature rises and decreases in humidity, to minimize mortality unrelated to treatment effects.

#### Hut collections

During the six days of exposure, hut baffles were closed each morning at 5 :30 am. Collectors equipped with headlamps carefully entered the huts and lowered the curtain separating the “hut” and the “veranda” compartments (**Fig. 1**). Mosquitoes were collected separately from each compartment using hemolysis tubes and were recorded according to their compartment of origin.

#### Tent collections

To collect mosquitoes inside the tents, the collector entered the net enclosure equipped with 15 cm X 15 cm holding cages and mouth aspirators. Upon entry, the 10 cm aperture was immediately closed to prevent mosquito escape. Mosquitoes were aspirated and transferred into cages labelled by night and cattle.

#### Transport and insectary procedures

Following collection, tubes and cages containing mosquitoes were covered with moistened mops to maintain humidity and transported within approximately 15 minutes in air-conditioned vehicle to the Diébougou insectary (insectary conditions: temperature 27 ± 2 °C; relative humidity 75 ± 5%; photoperiod 12:12 h light: dark).

#### Attractiveness measurements

Attractiveness was quantified as the number of mosquitoes attracted to cattle hosts and captured in huts or tents. Comparisons between treatment groups and sampling rounds were conducted to investigate whether ivermectin treatment modified host-associated odor cues, thereby influencing host attractiveness to mosquitoes and the resulting mosquito catches in huts and tents.

#### Mosquitocidal efficacy assessment

Mosquitocidal efficacy was defined as the mortality induced in wild mosquito populations after blood feeding on cattle hosts treated with mdc-STM-001 at two dose levels, compared with mosquitoes fed on untreated control cattle. Upon arrival at Diebougou insectary, a target of 100 fully engorged mosquitoes per tent collection cage was randomly selected, when sample size permitted, and transferred to squared holding cages for survival monitoring (15 cm X 15 cm). Mosquitoes collected in hemolysis tubes (hut collections) were transferred to mesh-covered holding cups (height × diameter = 7.6 × 6.3 cm), with 10 mosquitoes per cage. All mosquitoes were provided daily with a 10% glucose solution. Mosquito mortality was monitored daily for 14 consecutive days. Dead mosquitoes were counted daily, individually transferred to labelled 1.5-mL microcentrifuge tubes, and stored at -20 °C until taxonomic identification. Mosquitoes included in the survival monitoring subset were identified to genus, species complex, species group, or, when possible, species level using standard morphological identification keys [36] under a stereomicroscope (×40 magnification). All data were recorded in electronic spreadsheets (Microsoft Excel). Mosquitoes not selected for survival monitoring were frozen and subsequently identified to genus level.

### Ivermectin bioanalysis

Blood samples were planned to be collected from treated calves at each mosquito exposure time point and at additional predefined time-points (29 samples per animal) to characterize ivermectin pharmacokinetics (PK). For mdc-STM-001-treated animals in Groups 1-2 (dose 1.0 mg/kg) and 4-5 (dose 1.5 mg/kg), sampling included predose (prior to injection), after injection at 12h, 24h (Day After Injection, DAI 1), 36h, 48h (DAI 2), followed by one sample every 2 to 8 days until DAI 119. A final sample was collected immediately prior to explant excision, which occurred at different DAIs due to logistical constraints (DAI 135, 143 or 150). Control animals (Groups 3 and 6) were sampled at predose and on DAI 1, 15, 28, 56, 84 and 112 (7 samples per animal). Blood samples were gently agitated and stored on ice before centrifugation at +4°C (2000 rpm for 10 min) to isolate plasma, which was immediately stored at -20°C until shipment to the bioanalytical facility, where samples were stored at -80°C until analysis. Ivermectin plasma concentrations were quantified by liquid-liquid extraction using a qualified LC-MS/MS method including a low calibration range (0.2 - 20 ng/mL) and a high range (5 - 500 ng/mL) using K_2_-EDTA as anticoagulant and ivermectin-D_2_as internal standard. A mid calibration ranged from 1 to 100 ng/mL, derived from the high range, was also used. Quality control samples (0.6, 2, 10, 16 ng/mL low range; 3, 50, 80 ng/mL mid-range; 15, 250, 400 ng/mL high range) met the acceptance criteria (precision ≤ 15.00%, deviation ± 15.00%). Incurred sample reanalysis confirmed the ivermectin stability for at least 225 days when stored at -80°C. Plasma samples collected from cattle were stored for 70 - 220 days prior to analysis, supporting the validity of the ivermectin concentrations measured for pharmacokinetic analysis.

At the end of the study, control animals were returned to the herd, while treated animals were euthanized. On DAI 135 or DAI 143 or DAI 150 depending on the animal, injection sites and the associated depots with surrounding tissues were collected to quantify the remaining ivermectin and copolymers in the depot/tissue. These analyses were performed using ultra-performance liquid chromatography for ivermectin (limit of quantification: 0.9 µg/mL) and nuclear magnetic resonance spectroscopy for copolymers. Both analytical methods were developed for exploration purposes.

### Statistical analysis

All statistical analyses were conducted using the R software (version 4.5.1; R Core [37]). Except for survival data, models performance and goodness of fit were assessed using the “performance” package [38] and the DHARMa package for residual diagnostics [39].

Estimates were presented with their 95% confidence interval and statistical significance was considered at *p* < 0.05.

### Cattle follow up

Differences in weight gain between treatment groups were analyzed using the Kruskal-Wallis test and pairwise Wilcoxon rank sum exact test to compare treatments. Both tests were two sided. Confidence intervals (CI95%) were computed, and *p*-values were adjusted for multiple comparisons using the Holm’s correction method [40]. Analyses were performed on the 36 cattle (12 per treatment), irrespective of the hut of hut approach.

For coproscopic data, the statistical analysis is described in **supplementary file**.

### Pharmacokinetics

Pharmacokinetic parameters of ivermectin for all mdc-STM-001 treated groups at both doses and for both experimental approaches (hut and tent), were calculated over a 4-month period (*i.e.,* 119 days) using a non-compartmental analysis (Phoenix® WinNonlin® version 8.1; Certara USA, Inc., Princeton, NJ) performed on observed individual cattle concentrations.

The nominal times were used as no deviations from target sampling schedule were reported. These included the rate of systemic exposure (Cmax), the time to reach Cmax (Tmax), the extent of systemic exposure over 28 days (AUC0-28d), over 91 days (AUC0-91d) and over 119 days (AUC0-119d), concentrations at 28 DAI (C28d), 56 days (C56d), 91 DAI (C91d) and at 119 DAI (C119d). The apparent terminal half-life was not estimated due to the flat shape of the PK curves. The systemic exposure parameters were normalized to the actual administered dose calculated for each cattle using the weight of each syringe recorded before and after administration. The inter-individual variability in plasma levels and PK parameters was expressed as standard deviation (SD) in the tables and figures and was interpreted using the coefficient of variation (CV %) for each experimental group and approach (tent or hut). Daily individual cattle concentrations were used to refine the dose-response relationships; these daily concentrations were interpolated between observed data points using a nonparametric superposition approach, which does not rely on any predefined pharmacokinetic model.

### Effect of ivermectin on mosquito densities and diversity (in both tent and hut approaches)

For each experimental approach, we tested whether the number of mosquitoes captured per cattle was affected by ivermectin treatment and exposure round at the genus level. For the *Anopheles* genus specifically, we also examined whether mosquito abundance varied according to treatment, exposure round, and *Anopheles* species status. Generalized linear mixed models (GLMMs) were fitted using the *glmmTMB* package in R, assuming a negative binomial error distribution with quadratic parameterization to account for overdispersion in count data. Treatment, sampling round, and their interaction were included as fixed effects. For species-level analyses within the *Anopheles* genus, species identity and its interactions with treatment and sampling round were also considered as fixed effects. To account for repeated observations, cattle lot and cattle identity nested within cattle lot were included as random intercept effects. We further fitted a diversity model using the Shannon diversity index as the response variable to assess the effect of ivermectin treatment on species diversity within the *Anopheles* genus only. The Shannon diversity index was calculated using the *vegan* package [41].

Likelihood ratio tests were used to assess whether the inclusion of interaction terms and random effects improved model fit. Statistical inference for the final selected models was based on Type III analysis of deviance using Wald chi-square tests.

### Survival analysis

Due to insufficient survival sample sizes from the hut approach, survival analyses were performed using data from the tent traps only. Because the day of first survival monitoring actually corresponded to the second day post-blood feeding, two additional days were added to the time-to-event (death) variable prior to analysis. Kaplan-Meier survival curves [42] were generated and stratified by treatment group and genus, and for the Anopheles genus, by species.

The impact of treatment on mosquito survival was evaluated using mixed-effects Cox proportional hazards models [43], incorporating treatment and days post-injection (DAI) as interaction terms. Random effects included and individual bovine host identifier nested by batch, to account for intra- and inter-group variability.

Due to variability of mosquitoes fed on control animals, treatments effects were primarily assessed using 4-day cumulative mortalities. Therefore, an assay was excluded when mortality at 48h in control arm exceeded 20%. Analysis were also restricted to DAI with at least 3 calves in each arm The 4-day follow-up captured acute lethal effects of ivermectin ingestion, typically occurring within 2–3 days after feeding on treated blood, and has been already used to assess the mdc-STM-001 efficacy at the dose of 0.6 mg/kg [34]. Hazard ratios (4-day HRs) with their corresponding 95% confidence intervals (95% CI) were estimated, allowing quantification of the differential effect of treatment compared to the control group on mosquito mortality risk.

### Estimation of lethal concentrations and coverage duration

Dose–response relationships were modeled using the *drc* package in R [44], with 4-day cumulative mosquito mortality as the dependent variable, and the corresponding ivermectin plasma concentrations measured in treated cattle as the explanatory variable.

Cumulative 4-day mosquito mortality rates were calculated across the six exposed calves per night for each post-injection time point when mosquito exposure was performed. The corresponding ivermectin concentration was defined as the mean concentration across the six cattle.

### Ethical considerations

The study protocol has been approved by the institutional ethical committee with the reference number 25-2021/CEIRES/ 06/15/2021. Animal welfare requirements were applied following [45].

## Results

### Cattle follow up

A mild reaction (transient agitation) was observed in most animals following administration, irrespective of dose, treatment group or experimental approach (data not shown). No unscheduled deaths or adverse clinical signs were reported in any treatment group throughout the study, revealing the overall safety of the intervention. As expected with this formulation, localized swelling was the common reaction, occurring across treated animals. The data showed a favorable impact of ivermectin treatments on weight gain over the study period.

Quarterly weight intake (kg/90 days) showed a significant difference between control and treated groups (*p* = 0.032), with no significant uptake difference between treatment doses (pairwise comparison, Wilcoxon sum exact test, *p* = 0.838). The median (Q1, Q3) intake was 7 (1, 12) kg in the control group, compared to 16 (8, 24) kg and 15 (7, 23) kg in the mdc-STM-001 group at 1.0 mg/kg (*i.e.,* mdc-STM-1.0) and mdc-STM-001 group at 1.5 mg/kg (*i.e.,* mdc-STM-1.5), respectively. Detailed monthly weight measures for each group are provided in **supplementary file S1: Table 2**.

**Table 2.**
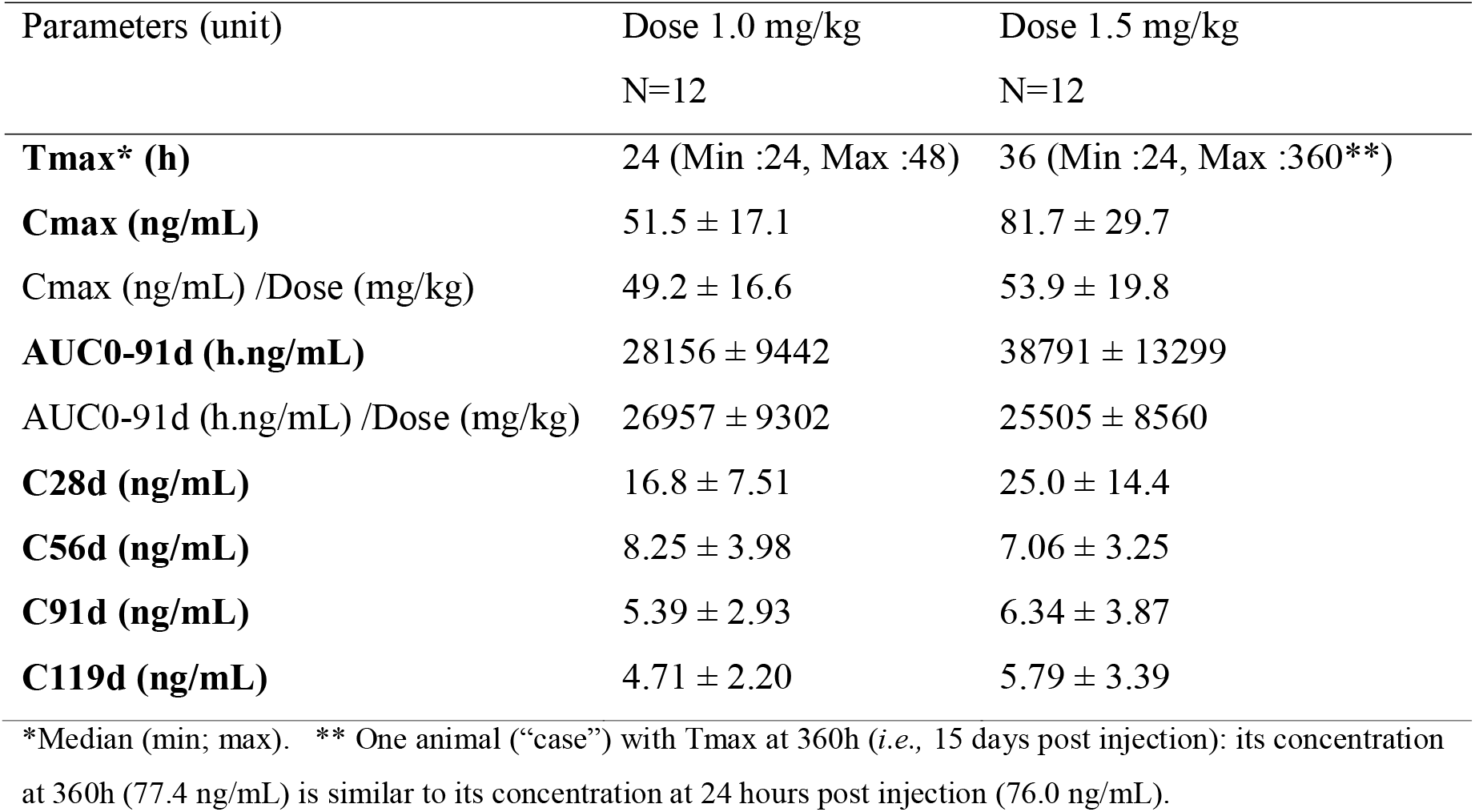
Main pharmacokinetic parameters (Mean ± SD) of ivermectin in cattle plasma after single subcutaneous administration of mdc-STM-001 at 1.0 and 1.5 mg/kg.

### Pharmacokinetic profile of ivermectin in the cattle plasma

A total of 760 blood samples were collected with 680 from mdc-STM-001 treated cattle (4 treated groups, 6 cattle per group, n=28 or 29 per cattle) and 80 from control cattle (2 control groups, 6 cattle per group, n= 6 or 7 per cattle). Concentration-time patterns were analyzed using ivermectin plasma concentrations measured until DAI 119 which is the last common sampling day for all animals. Ivermectin plasma concentrations were analyzed according to the exposure approach (*i.e.,* “hut” or “tent”) and to the dose level (*i.e.,* 1.0 or 1.5 mg ivermectin/kg). Individual concentration-time profiles of ivermectin after mdc-STM-001 injection are illustrated per exposure approach and per dose in **Fig. 2**. Ivermectin was not quantified in any of the samples collected in the control group during the study period and in any samples collected before ivermectin injection in the treated groups. After the subcutaneous injection of mdc-STM-001, ivermectin plasma concentrations were generally measured in all samples collected during the study, with quantifiable levels still present at DAI 112 days (for one animal only) and at DAI 119 and even at the last collection times (*i.e.,* DAIs 135 or 143 or 150 post-injection). As expected for a same dose level of mdc-STM-001, there were no significant differences in ivermectin plasma levels and its PK parameters between both exposure approaches, *i.e.,* “case” or “tent”, taking into account the inter-animal variability which generally tended to be similar or lower with “tent” compared to “case” (respectively, average CV% of plasma levels: 45% vs. 52% at 1.0 mg/kg and 55% vs. 57% at 1.5 mg/kg). Consequently, the overall PK interpretation was focused on the results obtained per dose level by merging the concentrations of both approaches (*i.e.,* n=12 animals per dose). To evaluate the mean PK profiles and the dose effect, an overlay of mean ivermectin plasma concentrations over time obtained at both doses is illustrated in **Fig. 3 A and B**. Additional figures, such as the individual kinetics illustrated in semi-log scale, the mean concentration-time profiles comparing both exposure approaches at a same dose (1.0 or 1.5 mg/kg) over the study duration and focused on the first 6 days (to better visualize the burst, *i.e.,* peak plasma level), are provided in the **supplementary file S1: fig. 1A, 1B and 1C**, respectively.

**Fig. 2.**
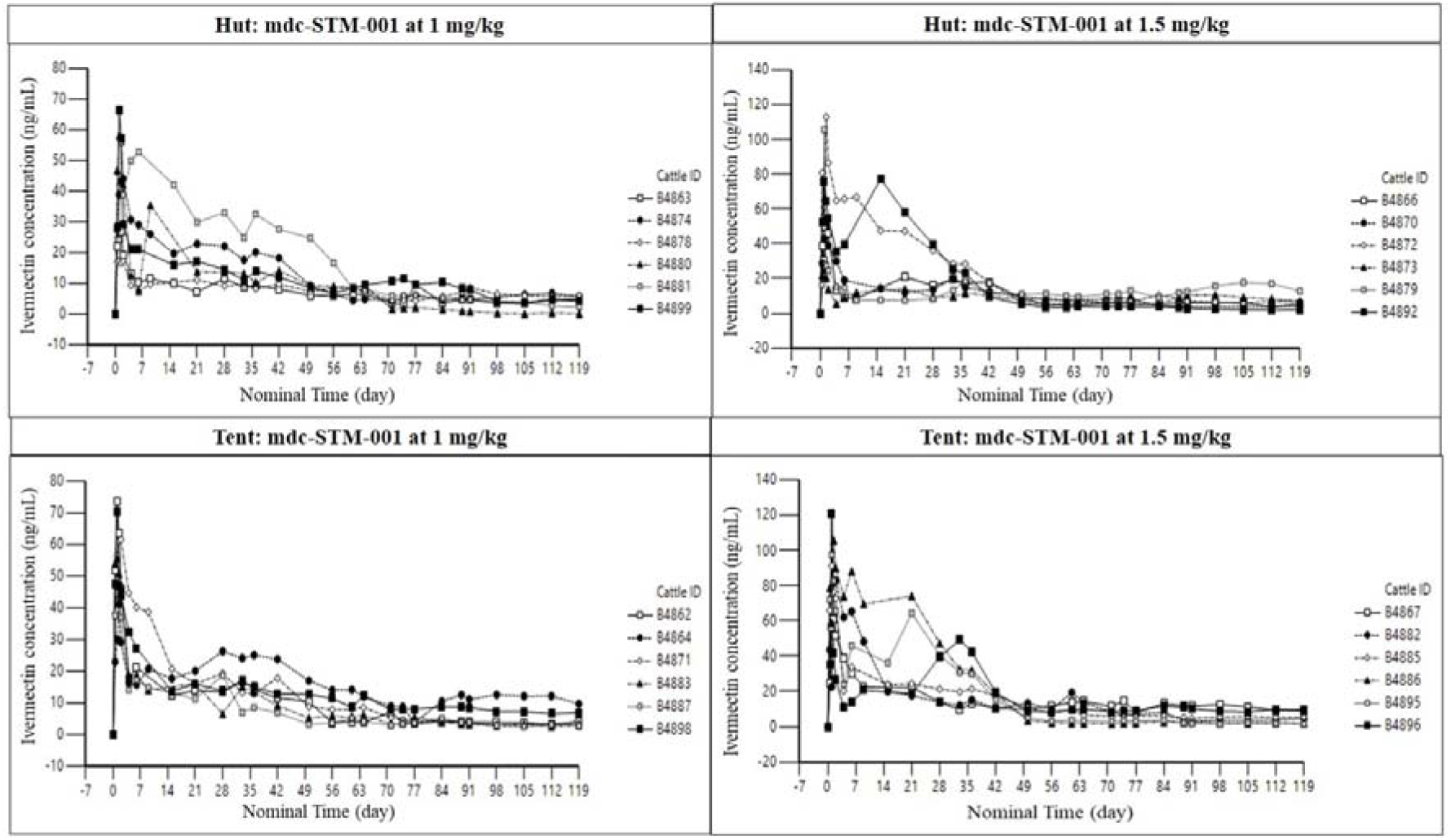
Individual kinetic profiles of ivermectin in cattle plasma after single subcutaneous administration of formulation (mdc-STM-001) per approach and per dose level (linear scale)

**Fig. 3.**
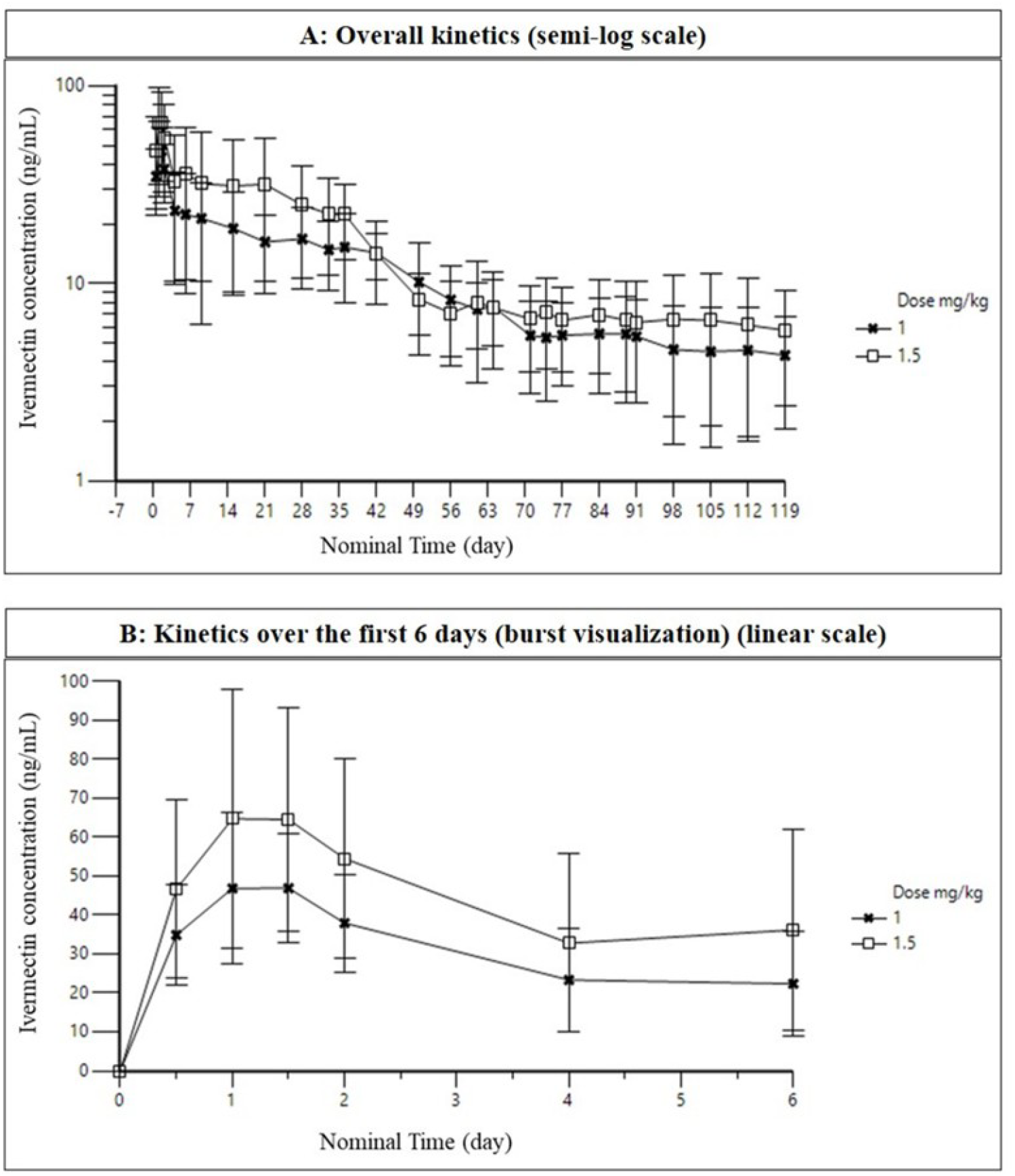
Overlay of mean (SD) kinetic profiles of ivermectin in cattle plasma after single subcutaneous administration of mdc-STM-001 at 1.0 and 1.5 mg/kg (N=12 per dose level). A: Overall kinetics in semi-log scale. B: Kinetics over the first 6 days for the burst visualization (linear scale).

After a single SC injection of mdc-STM-001, the shape of the PK curve was similar at both doses with an increase in systemic exposure when the dose increased from 1.0 to 1.5 mg/kg. The mean plasma levels of ivermectin increased to a maximal value reached at 24h for 1.0 mg/kg and at 36h for 1.5 mg/kg, then an approximate 2-fold fast decrease was observed until 4 days post-dose (at DAI 4: 23.3 ± 13.2 ng/mL and 32.8 ± 23.0 ng/mL, respectively).

Subsequently a slow release phase was observed until DAI 71 for 1.0 mg/kg (C71d: 5.4 ± 2.7 ng/mL) and until DAI 50 for 1.5 mg/kg (C50d: 8.3 ± 2.9 ng/mL). From these times, the plasma levels demonstrated a quite constant release rate of ivermectin, with mean ivermectin levels at 5.4 ng/mL for 1.0 mg/kg and at 6.3 ng/mL for 1.5 mg/kg at DAI 91 (*i.e.,* 3 months which is the target release duration). Beyond 3 months, the plasma levels still demonstrated a relatively constant release rate of ivermectin. The inter-variability (CV%) in ivermectin plasma levels was quite similar between both doses with on average 49% at 1.0 mg/kg and 56% at 1.5 mg/kg.

Considering 12 animals per dose level (*i.e.,* merged “case” and “tent” plasma data), the main pharmacokinetic parameters of ivermectin after a single SC injection of mdc-STM-001 are reported in **Table 2**. Additional PK parameters per dose level and those obtained for each exposure approach are given in the **supplementary file S1: Table 3 and Table 4**, **respectively**.

**Table 3.** Median survival times (days) for tent approach experimentation with their 95% confidence intervals.

| Species | Round | Control | mdc-STM-1.0 | mdc-STM-1.5 |
| --- | --- | --- | --- | --- |
| *All Anopheles* | 1 | 8 [7,8] | 3 [3,3] | 2 [2,3] |
|  | 2 | 9 [9,10] | 4 [4,5] | 3 [3,3] |
|  | 3 | 8 [8,9] | 6 [5,7] | 6 [6,7] |
|  | 4 | 7 [7,8] | 5 [5,6] | 7 [6,8] |
| *Anopheles coustani* | 1 | 7 [7,8] | 3 [3,3] | 3 [3,3] |
|  | 2 | 10 [9,10] | 5 [5,6] | 3 [3,3] |
|  | 3 | 8 [8,8] | 7 [6,8] | 7 [7,8] |
|  | 4 | 6 [6,7] | 5 [5,6] | 7 [7,8] |
| *Anopheles funestus* | 1 | 5 [4,9] | 2 [2,2] | 2 [2,2] |
|  | 2 | 9 [7,14] | 3 [2,3] | 2 [2,2] |
|  | 3 | 8 [7,9] | 4 [4,8] | 3 [3,4] |
|  | 4 | 9 [7,10] | 6 [5,10] | 5 [5,8] |
| *Anopheles rufipes* | 1 | 9 [7,10] | 2 [2,2] | 2 [2,2] |
|  | 2 | 11 [9,13] | 3 [3,3] | 2.5 [2,3] |
|  | 3 | 9 [9,10] | 3 [3,5] | 5 [4,6] |
|  | 4 | 8 [7,9] | 4 [3,5] | 6 [5,8] |
| *Anopheles squamosus* | 1 | 8 [5,13] | 3 [3,3] | 2 [2,2] |
|  | 2 | 8 [8,9] | 4 [3,4] | 3 [3,3] |
|  | 3 | 9 [7,10] | 7.5 [6,9] | 8 [6,10] |
|  | 4 | 9 [7,10] | 7 [5,9] | 8 [5, NA] |

**Table 4.** Ivermectin concentrations inducing 50% lethal concentrations (LC) for *Anopheles* at DAI4 for both doses. Mean concentrations (ng/mL) are given with lower and upper confidence intervals (95% CI)

| Species | LC50 (ng/ml) for mdc-STM-1.0 | LC50 (ng/ml) for mdc-STM-1.5 |
| --- | --- | --- |
| All *Anopheles* | 14.2 [13.4-15] | 9.4 [8.8-10] |
| *An. coustani* | 19.2 [18.1-20.3] | 12.4 [11.6-13.2] |
| *An. funestus* | 6.9 [5.0-9.4] | 5.1 [3.9-6.6] |
| *An. rufipes* | 5.7 [4.3-7.6] | 5.8 [4.6-7.2] |
| *An. squamosus* | 14.1 [12.3-16.1] | 8 [6.6-9.8] |

After a single SC injection of mdc-STM-001 at 1.0 mg/kg, the mean peak plasma level of ivermectin (*i.e.,* Cmax or burst) at 51.5 ± 17.1 ng/mL was reached at a median Tmax of 24 hours post-dose. The Cmax inter-variability (CV%) was moderate at 33% with 22% for “tent” and 43% for “case”. Over 91 days post-dose (*i.e.,* target duration of 3 months), the mean ivermectin extent of exposure (*i.e.,* AUC) at 28156 ± 9442 ng.h/mL represented 90% of the AUC over 119 days (*i.e.,* 4 months), showing that most of the release of ivermectin from the SC depot lasted up to 3 months, accompanied by a moderate AUC inter-variability of 34% with 25% for “tent” and 42% for “case”. The mean concentration at Day 91 was 5.4 ± 2.9 ng/mL (CV%: 54%) with a quite similar inter-variability between “tent” (60%) and “case” (53%). After single SC injection of mdc-STM-001 at 1.5 mg/kg, the mean peak plasma level of ivermectin at 81.7 ± 29.7 ng/mL was reached at a median Tmax of 36 hours post-dose. The Cmax inter-variability was moderate at 36% with 22% for “tent” and 50% for “case”. Over 91 days post-dose, the mean Ivermectin extent of exposure at 38791 ± 13299 ng.h/mL represented also 90% of AUC119d, accompanied by a moderate inter-variability of 34% with 24% for “tent” and 44% for “case”. The mean concentration at Day 91 (*i.e.,* target duration of ivermectin release) was 6.3 ± 3.9 ng/mL (CV%: 61%) with 75% for “tent” and 53% for “case”. For both doses, the mean concentrations after each month post subcutaneous injection (C28d, C56d, C91d, C119d) showed a slow decrease from DAI 56 to DAI 119, remaining between approximately 5 and 8 ng/mL with a moderate to high inter-variability ranging from 54 % to 61%. Over the dose range from 1.0 to 1.5 mg/kg, the systemic rate and extent of exposure of ivermectin increased proportionally with the dose increase, as Cmax/Dose and AUCs/Dose remained quite similar between both doses (high to low dose ratios from 0.9 to 1.1). The analysis of each animal’s depot collected at the end of the *in vivo* phase*, i.e.,* Day 135 or Day 143 or Day 150 after a single subcutaneous administration, showed that the average percentage of remaining amount of ivermectin not released from the depot was around 18% of the injected amount except for Group 1 (7%, hut 1.0 mg/kg).(**supplementary file S1: Table 5**).

**Table 5.** Time period during which the formulation administered at two doses reaches ivermectin plasma concentration values above lethal concentrations inducing 50% mortalities, over a 4-day period. The time period (day) corresponding to mean LC50 values are given and those corresponding to lower and upper confidence intervals (95%), respectively, are given in brackets.

| Treatment | ID bovin | All *Anopheles* | *An. coustani* | *An. funestus* | *An. rufipes* | *An. squamosus* |
| --- | --- | --- | --- | --- | --- | --- |
| mdc-STM-1.0 | B4862 | 35 [35-37] | 8 [6-9] | 71 [50-72] | 72 [52-73] | 36 [34-40] |
|  | B4864 | 55 [53-62] | 47 [46-48] | 119 [119-119] | 119 [119-119] | 61 [51-101] |
|  | B4871 | 45 [44-45] | 28 [15-28] | 67 [49-71] | 69 [65-75] | 45 [43-46] |
|  | B4883 | 34 [33-35] | 3 [3-6] | 46 [41-79] | 64 [45-82] | 34 [7-37] |
|  | B4887 | 29 [29-30] | 3 [3-28] | 41 [31-84] | 44 [39-86] | 29 [29-30] |
|  | B4898 | 38 [35-40] | 11 [10-12] | 108 [68-119] | 119 [96-119] | 38 [33-51] |
|  | Total | 39.3 [38.2-41.5] | 16.7 [13.8-21.8] | 75.3 [59.7-90.7] | 81.2 [69.3-92.3] | 40.5 [32.8-50.8] |
| mdc-STM-1.5 | B4867 | 74 [27-74] | 24 [23-25] | 119 [109-119] | 119 [106-119] | 101 [74-110] |
|  | B4882 | 63 [63-64] | 61 [15-62] | 84 [68-86] | 70 [67-85] | 66 [63-68] |
|  | B4885 | 46 [45-47] | 40 [38-42] | 62 [51-106] | 61 [49-63] | 48 [45-51] |
|  | B4886 | 44 [43-44] | 41 [41-42] | 47 [46-48] | 47 [45-48] | 45 [43-46] |
|  | B4895 | 42 [42-43] | 40 [40-41] | 48 [46-49] | 47 [45-48] | 44 [42-46] |
|  | B4895 | 45 [45-46] | 42 [41-43] | 119 [92-119] | 119 [89-119] | 47 [45-92] |
|  | Total | 52.3 [44.2-53] | 41.3 [33-42.5] | 79.8 [68.7-87.8] | 77.2 [66.8-80.3] | 58.5 [52-68.8] |

### Ivermectin effect on mosquito abundance and diversity

A total of 34,249 mosquitoes were collected using tent traps and 2,145 from experimental huts over the study period, with notable variations in genus composition between the two sampling methods (**Fig. 4**). Overall, *Anopheles* was the predominant genus particularly in tent sampling approach accounting for 52.67% of specimens collected and 35.37% in experimental huts. *Culex* represented the second most abundant genus, comprising 35.03% of the tent trap collections and 44.13% of those from huts. *Mansonia* contributed 11.17% and 6.94% of captures in tent traps and huts, respectively. In contrast, *Aedes* was relatively scarce in tent traps (1.13%) but appeared proportionally higher in huts (13.56%).

**Fig. 4.**
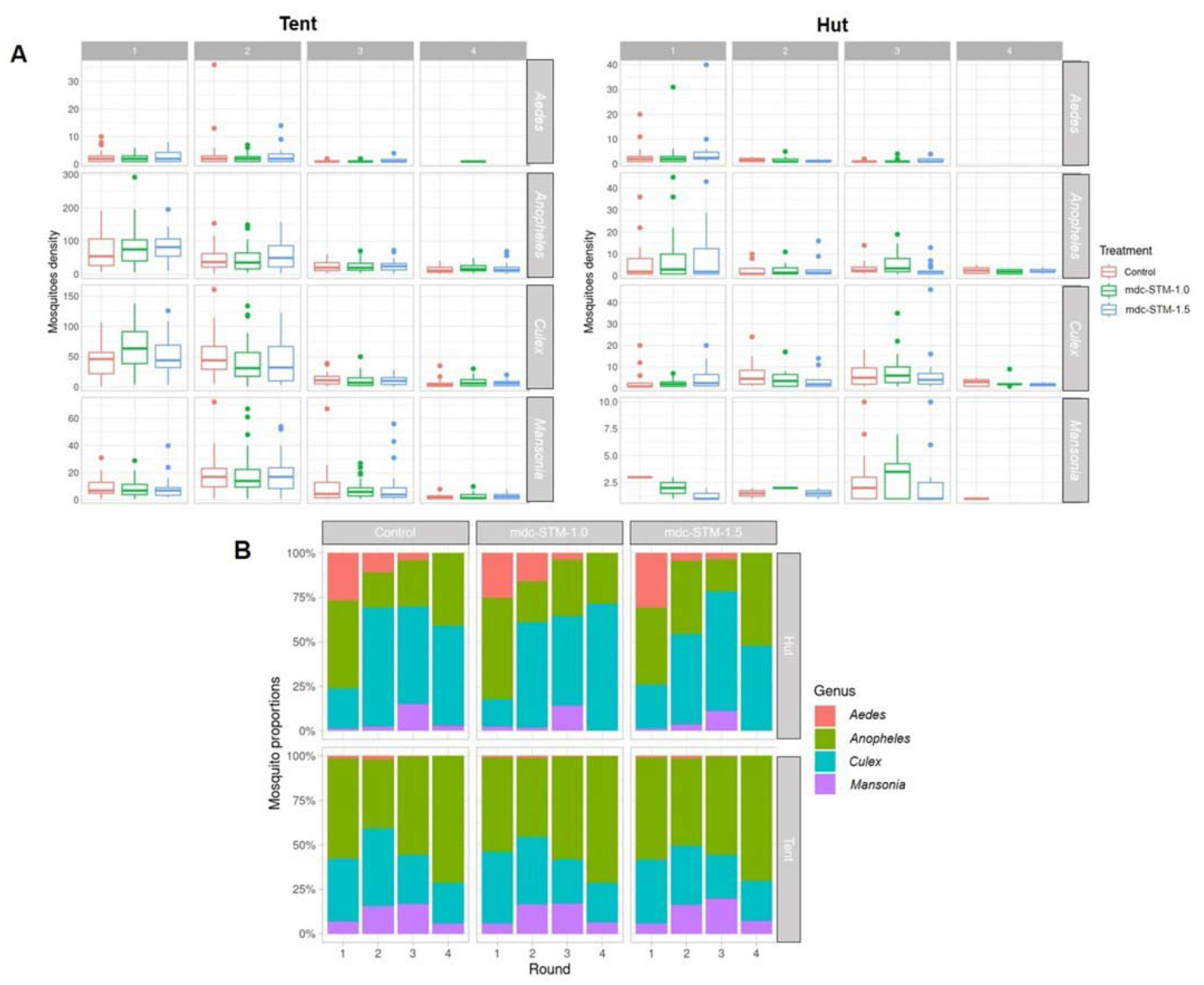
Overall mosquitoes catches density and genus composition by round, treatment, and experimental approach. A. Mosquito catch density, expressed as the number of mosquitoes collected per round. B. Relative abundance of mosquito genera, expressed as the proportion of the total catch represented by each genus for each round, treatment, and experimental approach.

While both methods sampled similar mosquito genera, tent traps yielded a substantially higher number of mosquitoes.

Based on the log-likelihood, the model including both interaction terms and the random effect was selected as this model was the best describing the data. The ivermectin treatment had no significant effect on overall mosquito densities under either sampling approach (treatment effect: tent approach, LRT χ² = 0.84, *p* = 0.65; hut approach, LRT χ² = 0.11, *p* = 0.94; see **supplementary file S1: Table 6 a and b**, respectively).

Similarly, neither the treatment × genus nor the treatment × round interactions were significant. In contrast, mosquito densities varied significantly among genera and across exposure rounds. The significant genus × round interaction further indicated that temporal changes in mosquito density differed among genera (for tent approach, LRT χ²_9_ = 155.14, *p* < 0.001, for hut approach, LRT χ²_9_ = 41.89, *p* < 0.001).

In tent catches, 10 species were identified among the *Anopheles* genus while only 8 were identified in huts, with *An. coustani* being by far the dominant species in both approaches (see F**ig. 5**). In tents, the second most abundant Anopheles species was *An. rufipes*, followed by *An. squamosus* and *An. funestus*. *An. gambiae s.l.* was the 5^th^ most present mosquito, with densities per cattle night being < 5. In the huts, this species represented the second most abundant mosquitoes. The relative abundance and exact numbers of the different species of mosquitoes from the other genera trapped are given in the **supplementary file S1:** F**ig. 2**. The *Culex* genus exhibited notable diversity with 13 species recorded in tents. Its diversity decreased in huts, with only 8 species recorded. *C. univittatus*, *C. quinquefasciatus*, and *C. robinotus* were among the most frequently recorded species. *Aedes* genus showed similar diversity in tents with 13 species identified. In huts, *Aedes* diversity decreased greatly, with only 3 species identified there. *Ae. vittatus* was by far the most abundant *Aedes* species. *Mansonia uniformis* dominated within the *Mansonia* genus, accounting for more than half of its total captures in both approaches.

**Fig. 5.**
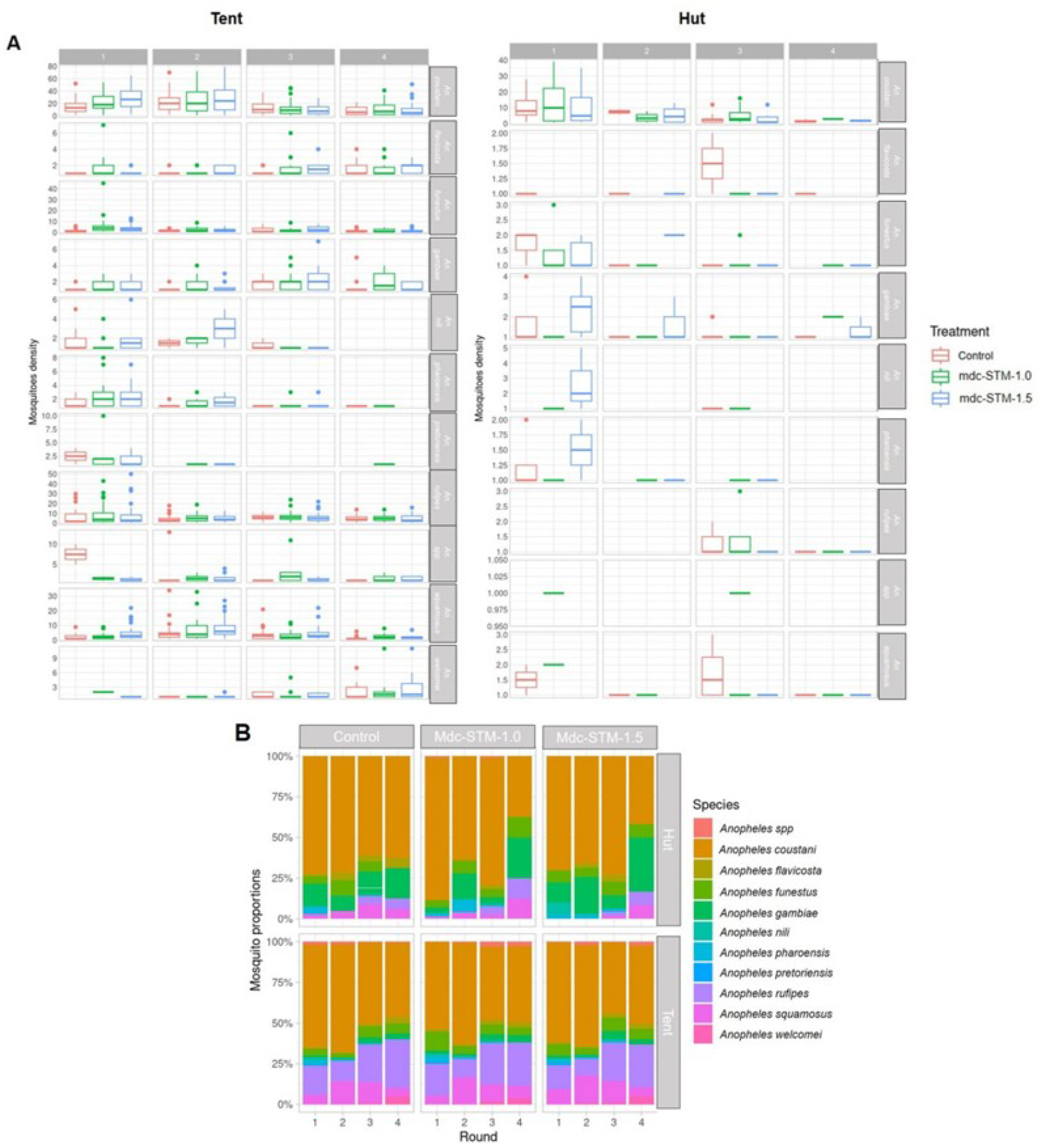
Mosquitoes catch density and species composition of *Anopheles* mosquitoes by round, treatment, and experimental approach. A. Mosquito catch density for each *Anopheles* species, expressed as the number of mosquitoes collected per round. B. Relative abundance of *Anopheles* species, expressed as the proportion of the total catch represented by species.

When testing treatment effect among *Anopheles* species, it appears that overall the treatment did not have a significant effect on mosquitoes’ densities for both tent and hut approaches. The treatment x species was not significant either. However, post-hoc procedure showed that densities were significantly different between treatments for *An. coustani*, *An. funestus* and *An. squamosus* during the first round of exposure only for the tent approach (**see supplementary file S1: Table 7**).

The Shannon diversity index revealed relatively stable *Anopheles* community diversity across sampling rounds and treatment groups (**Fig. 6)**. In experimental huts, diversity values generally remained low to moderate, with most Shannon index values below 1. Although slight increases were observed in the ivermectin-treated groups during round 1 sampling particularly, these differences were not consistently maintained over time. In contrast, collections from tent approach exhibited higher Shannon diversity values overall, ranging between 0.8 and 1.5 across all treatment groups and sampling rounds.

**Fig. 6.**
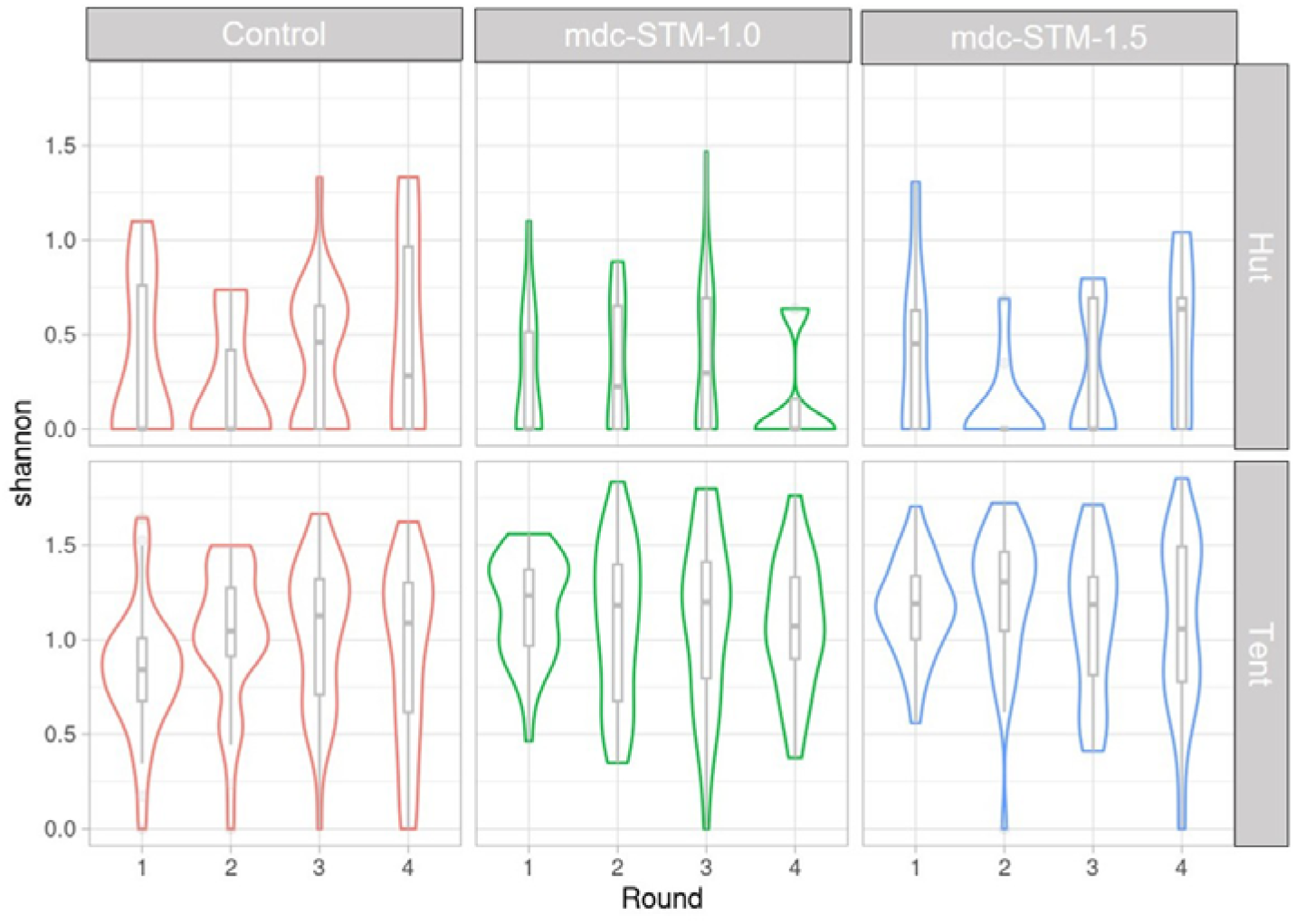
Distribution of Shannon’s index for *Anopheles* mosquitoes.

Temporal fluctuations in diversity were evident in both collection settings, as illustrated by changes in the distribution and spread of Shannon index values among rounds. However, the magnitude of these fluctuations was comparable between treated and control groups, and no persistent shift in community diversity was observed following treatment.

### Mosquitocidal activity evaluation

A total of 12,330 *Anopheles* specimens from the tent trap approach were monitored for survival; 3281 in control cattle, 4356 in cattle treated with 1.0 mg/kg IVM, and 4693 in cattle treated with IVM dose 1.5 mg/kg. Analyses were conducted for all *Anopheles* species combined and for the following species individually: *An. coustani*, *An. rufipes*, *An. funestus*, and *An. squamosus*. Survival models for the remaining trapped species, including *An. gambiae* s.l., did not converge because of small sample sizes. Therefore, these species were excluded from the analysis. All DAIs for which mortality exceeded 20% at 48 h of survival observations in the control group were excluded from further analysis.

### Mortality rates

The cumulative mortality rates of *Anopheles* mosquitoes using tent-based approach were evaluated across the four collection rounds to assess the effect of the ivermectin slow-release formulation at both doses. The analysis revealed species-specific responses to ivermectin exposure (**Fig. 7)**. Across all *Anopheles* mosquitoes combined, mortality remained low and relatively stable in the control group, globally below 20%, whereas both ivermectin-treated groups exhibited substantially higher mortality, particularly during the first 40 days after injection (DAI). Mortality gradually declined thereafter but remained above control levels.

**Fig. 7.**
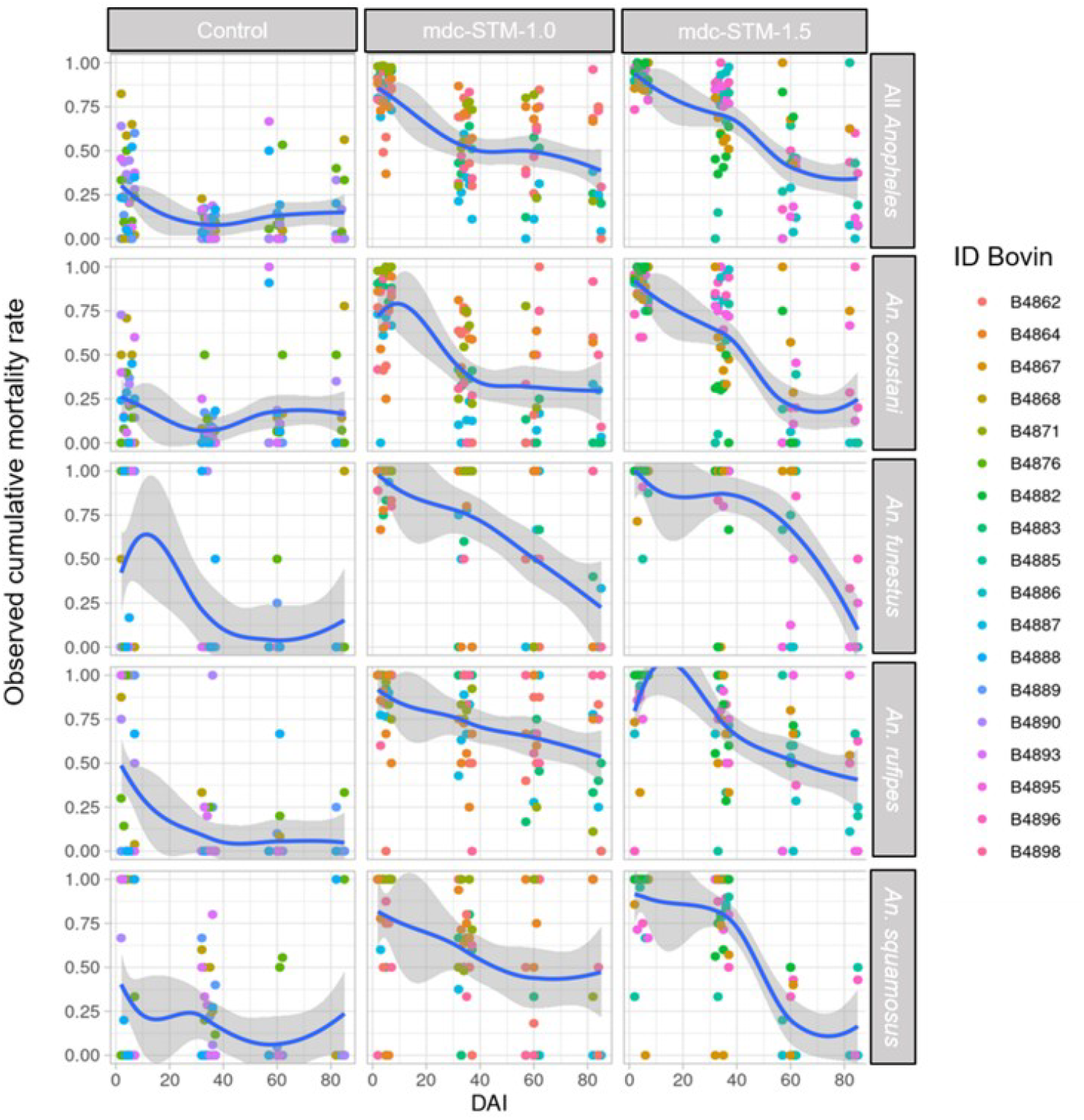
Cumulative mortality rates of *Anopheles* mosquitoes for the 4-day periods post exposure using tent-based approach on cattle treated with ivermectin long-acting formulation at the doses 1.0 mg/kg (mdc-STM-1.0) and 1.5mg/kg (mdc-STM-1.5) over time (days after injection, DAI). Cumulative mortality rates were assessed for all *Anopheles* species combined and for the most abundant species. Each point corresponds to the cumulative mortality recorded from mosquitoes associated with an individual bovine (identified by color). Blue lines indicate LOESS (Locally Estimated Scatterplot Smoothing) regression fits with 95% confidence intervals (grey area). Mortality remained low in the control group, whereas both ivermectin-treated groups showed elevated mortality during the early post-treatment period followed by a progressive decline over time. The mdc-STM-1.5 globally maintained higher mortality rates for a longer duration than mdc-STM-1.0.

The 1.5 mg/kg dose (mdc-STM-1.5) globally maintained slightly higher mortality (60-65%) than the 1.0 mg/kg dose (50-55%) during the later stages of follow-up.

For *Anopheles coustani,* mortality patterns closely look like those observed for the overall mosquito population. Mortality was highest shortly after treatment administration, reaching approximately 70-80% in both ivermectin-treated groups, before progressively declining with increasing DAI. A pronounced treatment effect was also observed for *Anopheles funestus*. In both ivermectin-treated groups mortality remained very high during the early post-treatment period (approximately 90-100% within the first 10-20 DAI) and declined steadily over time. The decline appeared particularly pronounced beyond 50-60 DAI, although mortality levels generally remained above those observed in the control group. Similarly, *Anopheles rufipes* showed elevated mortality in treated cattle groups compared with controls. Mortality remained high during the first month following treatment (approximately 75-90%) and subsequently declined, although the decrease was more gradual than that observed for *An. funestus.* The 1.5 mg/kg treatment appeared to maintain higher mortality over time.

For *Anopheles squamosus*, mortality patterns also reflected a treatment effect. Both doses induced high mortality during the early sampling period with approximately 80-85% within the first 10-20 DAI. A marked decline was observed after approximately 40-50 DAI, especially in the 1.5 mg/kg dose group.

### Survival and Hazard Ratios

Following treatment, both mdc-STM-1.0 (IVM dose of 1.0 mg/kg) and mdc-STM-1.5 (IVM dose of 1.5 mg/kg) induced a marked mosquitocidal effect in tent-based approach, with *Anopheles* survival probability significantly reduced compared to the control group (**Fig. 8**). Kaplan-Meier survival analysis showed that exposure to ivermectin-treated cattle reduced mosquito survival. Although the magnitude of the effect declined over time, both doses allowed mosquito mortality at a greater rate than in the control group. Hazard ratios (HRs) derived from Cox proportional hazards models quantified the treatment effects over 4-day intervals (**Fig. 9**). Both doses achieved HRs well above the WHO-recommended threshold of 4 for endectocidal efficacy against malaria vectors during early rounds. At 4 days post-treatment, mdc-STM-1.0 exhibited HRs ranging from 4.97 to 8.98 across rounds, while mdc-STM-1.5 achieved HRs between 2.84 to 16.40, underscoring the potent initial impact of both doses.

**Fig. 8.**
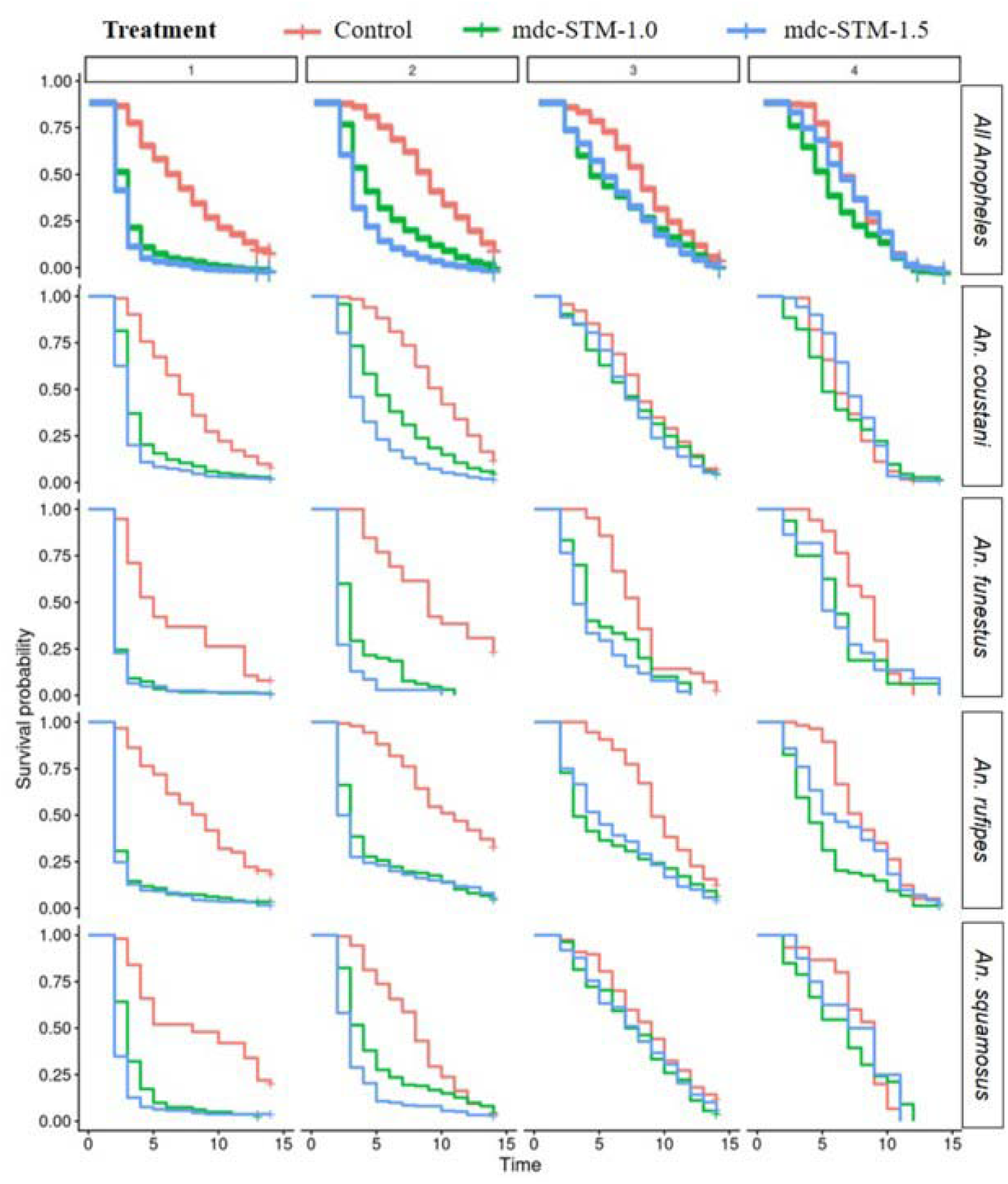
Survival of *Anopheles* mosquitoes by treatment and collection round using tent-based approach. Kaplan-Meier survival curves over two weeks follow up post-blood meal for mosquitoes fed on control cattle (red) and ivermectin-treated cattle (mdc-STM-1.0: green; mdc-STM-1.5: blue) across four rounds. Both ivermectin treatments significantly reduced survival compared to controls, with the strongest effects observed in early rounds (*p* < 0.001).

**Fig. 9.**
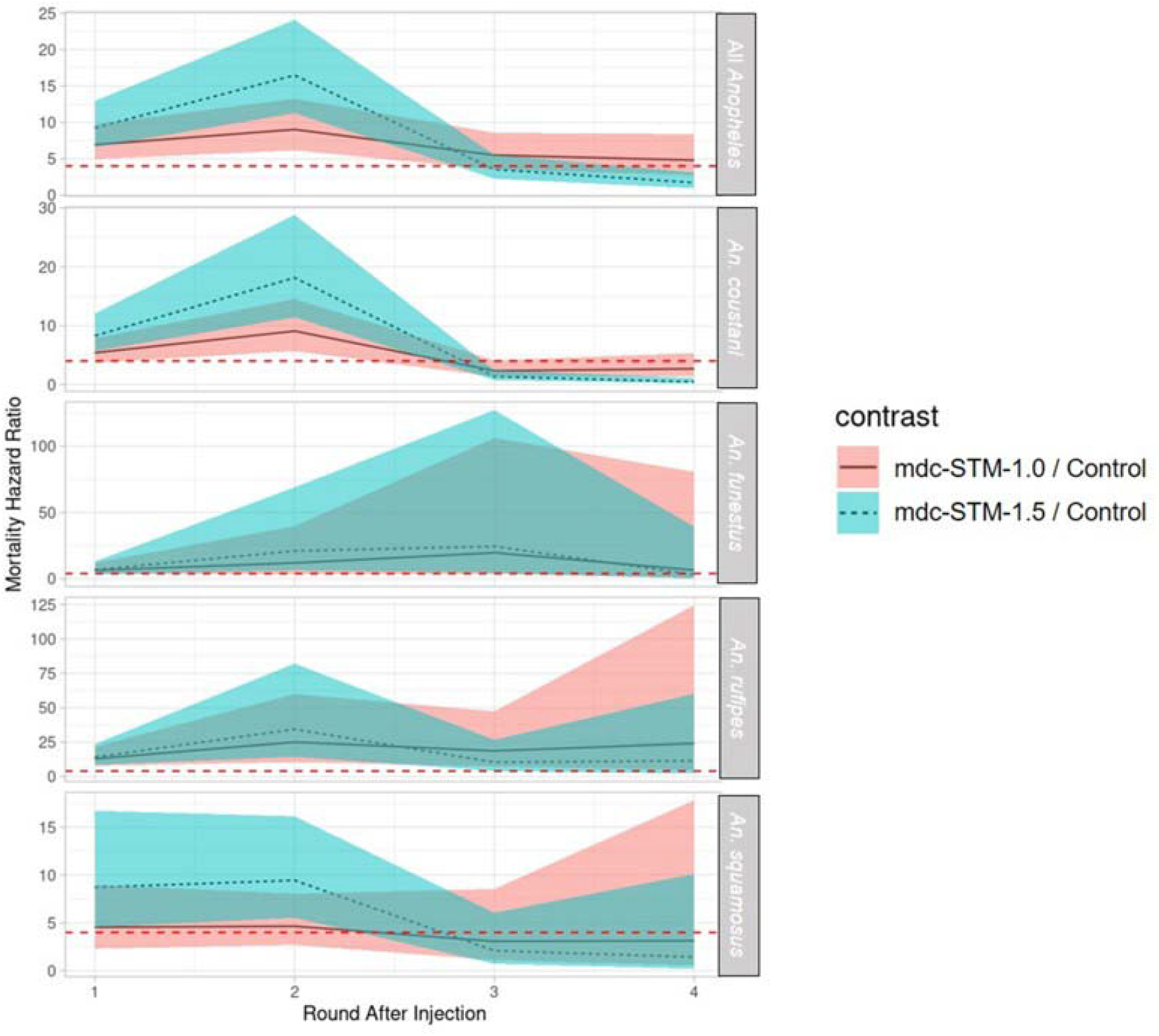
Mortality Hazard Ratios (HRs) of *Anopheles* mosquitoes. Panels show the aggregated data for all species pooled together (All *Anopheles*) followed by species-specific breakdowns for most abundant species (*An. coustani, An. funestus, An. rufipes, An. squamosus*). Contrasts show mortality risk relative to the control group for mdc-STM-1.0 (dose 1.0 mg/kg of the ivermectin formulation) and mdc-STM-1.5 (dose 1.5 mg/kg of the ivermectin formulation) ; shaded regions represent the corresponding 95% confidence intervals. The horizontal red dashed line marks the WHO-recommended threshold value of HR > 4 required to establish significant endectocidal efficacy against malaria vectors.

There was a significant overall treatment effect, which varied according to *Anopheles* species and study round (treatment effect: LRT χ² = 164.13, *p* < 0.001; treatment × species interaction: LRT χ² = 33.00, *p* < 0.001; treatment × round interaction: LRT χ² = 164.73, *p* < 0.001). Interestingly, regardless of treatment, there was also a significant interaction between round and species (LRT χ² = 28.34, *p* = 0.0008), indicating that overall mortality varied among rounds in a species-specific manner.

### Median survival times

The analysis of median survival times revealed a variation between treatment groups and sampling rounds (**Table 3**). Across all *Anopheles* species combined, those from the control group presented median longevity ranging from 7 to 9 days. In contrast, survival was reduced in both ivermectin-treated groups during the first two sampling rounds, reaching median values of 2-4 days. Although survival gradually increased in later rounds, it remained below the control levels. Among species, median survival for *Anopheles coustani* declined from 7-10 days in the control group to 3-5 days in treated groups during the first two rounds. Survival increased during rounds 3 and 4, approaching control values.

A particularly pronounced reduction was observed in *Anopheles funestus*. Median survival in the control group remained (around 9 days) whereas mosquitoes exposed to treated cattle survived 2-3 days during the first two rounds. Although survival increased during later rounds, median longevity remained lower than in controls.

*Anopheles rufipes* also showed strong treatment-associated reductions in survival. Median longevity decreased from 8-11 days in control to 2-3 days during rounds 1 and 2 in both treated groups. For *Anopheles squamosus,* treatment effects were also evident during the early post-treatment period. Median survival declined from approximately 8-9 days in controls to 2-4 days in treated groups during rounds 1 and 2.

### Lethal concentration and coverage duration

Dose-response relationships were established for *Anopheles* collected in tent-based approach and for both doses of the formulation (mdc-STM-1.0 and mdc-STM-1.5) injected to cattle, using cumulative mortality recorded at 4 days post-exposure (**Fig. 10)**. Across all panels, mortality increased with increasing ivermectin concentration. The fitted dose-response curves show that mortality rose at low to intermediate ivermectin concentrations before reaching a plateau close to 100% indicating a clear concentration-dependent effect of the formulation.

**Fig. 10.**
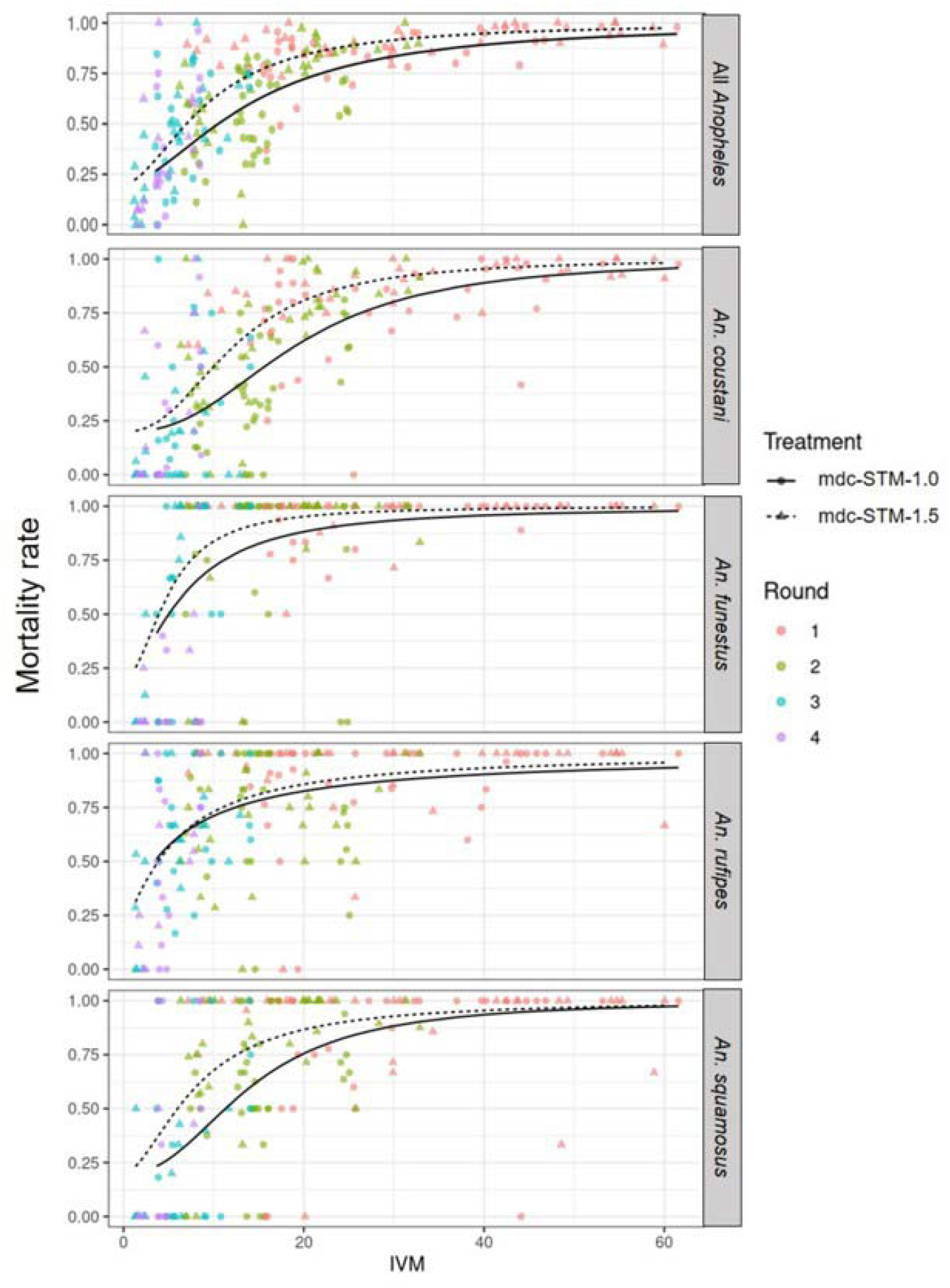
Dose-response relationship curves of *Anopheles* mosquitoes for mortality rates at 4 days. The panels illustrate the proportion of affected mosquitoes (y-axis) relative to ivermectin plasmatic concentration (x-axis) for all *Anopheles* species combined and stratified for abundant species (*An. coustani, An. funestus, An. rufipes* and *An. squamosus*. Points represent observed cumulative mortality and line types represent fitted dose-response curves for cattle treated with mdc-STM-1.0 (solid line) and mdc-STM-1.5 (dashed line)

Model selection analyses demonstrated that incorporating formulation dose as an explanatory variable significantly improved model fit compared with models excluding it. Dose-inclusive models globally produced lower AIC and BIC values and higher log-likelihoods than models without dose (**supplementary file S1: Table 8.)**, hence the inclusion of formulation dose in subsequent dose-response analyses. Species-specific differences were evident in susceptibility to ivermectin for both doses. *An. funestus* was the most susceptible species achieving high mortality at relatively low ivermectin concentrations. This high sensitivity was reflected by the lowest estimated LC50 (**Table 4**). *Anopheles rufipes* displayed a similarly pronounced response with mortality increasing rapidly across the concentration range and remaining high even at moderate ivermectin levels. In contrast *An. coustani* exhibited the lowest susceptibility. Mortality increased more gradually with concentration and higher ivermectin levels were required to achieve equivalent mortality compared with the other species.

Enhanced efficacy of the higher dose was observed for *An. coustani* as evidenced the comparison of LC50 estimates between doses that showed that the LC50 of mdc-STM-1.0 was significantly higher than that of mdc-STM-1.5 (**supplementary file S1. Table.9**, LC50 ratio = 1.55, *p <* 0.001). *An. squamosus* displayed an intermediate susceptibility profile.

Importantly, *An. coustani* and *An. squamosus* were the species for which a significant difference in LC50 was detected between doses with the LC50 of mdc-STM-1.0 being respectively 1.55 and 1.76-fold higher than that of mdc-STM-1.5 (LC50 ratio = 1.55 for *An. coustani* and 1.76 for *An. squamosus*, *p <* 0.001). In contrast, for *An. rufipes* and *An. funestus* no significant differences in LC50 were observed between doses (*p* > 0.05 for these species). **Table 5** summarizes the time intervals during which the two ivermectin doses allowed maintaining plasma concentrations above the lethal thresholds required to induce 50% mortality (LC50). For the pooled *Anopheles,* the LC50 concentrations were maintained on average for 39.3 days for mdc-STM-1.0 and 52.3 days for mdc-STM-1.5. Species-specific differences were observed in the duration of LC50 coverage. *Anopheles coustani* exhibited the shortest period of susceptibility, with LC50 plasma concentrations remaining globally for 16.7 and 41.3 days respectively for mdc-STM-1.0 and mdc-STM-1.5. In contrast, *Anopheles funestus* showed the longest duration of LC50 coverage among species examined. LC50 plasma concentrations remained for an estimated 75.3 days for mdc-STM-1.0 and 79.8 for mdc-STM-1.5. Similarly, *Anopheles rufipes* maintained LC50 coverage for extended periods reaching respectively 81.2 and 77.2 days for mdc-STM-1.0 and mdc-STM-1.5. *Anopheles squamosus* displayed an intermediate response. Its LC50 plasma concentration was sustained on average for 40.5 days for mdc-STM-1.0 and 58.5 days for mdc-STM-1.5. Although some inter-animal variability was observed, treated cattle maintained plasma concentrations above the LC50 threshold for several weeks following a single injection of the formulation.

Overall, *Anopheles funestus* and *Anopheles rufipes* were the most responsive species, maintaining lethal ivermectin exposure for the longest periods.

## Discussion

Beyond IVM well-documented effects against parasitic infections, its mosquitocidal properties have been increasingly explored as a potential vector control strategy [46, 47]. Recent clinical and entomological trials have provided encouraging evidence that ivermectin can reduce malaria transmission by increasing mortality in blood-feeding mosquitoes [48]. However, while some studies reported reductions in malaria incidence [18, 19], a trial by Somé *et al.* has pointed out the short-term effects of ivermectin MDA campaign which may limit the overall efficacy of such intervention [22]. The findings of all these studies highlight the challenge of translating short-lived mosquitocidal effects into sustained community-level impact and stimulate the interest in long-acting formulations capable of maintaining effective mosquitocidal systemic concentrations for extended periods after a single injection. Among these, BEPO^®^ technology has emerged as a promising controlled-release platform. This in situ forming depot gradually releases ivermectin over several weeks to months, thereby overcoming the pharmacokinetic limitations of conventional formulations [33, 49, 50].

Studies in cattle models have demonstrated that the BEPO^®^ based long-acting injectable ivermectin formulations can maintain plasma concentrations within the mosquitocidal range for several months and affect mosquito survival, especially *An. coluzzii* under experimental conditions [34, 50]. And model-based projections suggested that a single campaign using long-acting injectable ivermectin could reduce malaria transmission by 16-29%, with the impact increasing to 33-57%, 48-74% and 59-78% after respectively two, three and four consecutive campaigns [51].Our study was designed as a translational step toward the development of long-acting ivermectin formulations for malaria vector control. By evaluating a BEPO^®^ - based formulation under semi-field conditions in cattle, we generated critical pharmacokinetic and entomological data to support the further development of this approach for human use as a complementary strategy to reduce malaria transmission. The trial was established to target malaria vectors naturally attracted to treated cattle, with a focus on zoophilic and secondary vectors species that are often underrepresented in conventional studies. While members of *Anopheles gambiae* complex were detected, mosquito populations were predominantly composed of secondary vectors including *An. funestus, An. coustani, An. rufipes* and *An. squamosus.* This is particularly relevant given the growing recognition of the contribution of secondary vectors to residual malaria transmission.

### Pharmacokinetic and pharmacodynamic profile of the ivermectin formulation mdc-STM-001

The mdc-STM-001 long-acting formulation was tolerated at both 1.0 and 1.5 mg/kg in the young male cattle whatever the mosquito’s exposure approach (*i.e.,* tent or hut) with no observable adverse effects. Based on pharmacokinetics and efficacy results, this formulation is confirmed to be a promising lead formulation for further development toward human use. In cattle, pharmacokinetic results showed that ivermectin was delivered into bloodstream slowly, reaching a mean concentration of 8 ng/mL and 6 ng/mL respectively after 2 and 3 months post injection which is above the target duration for malaria indication [52], with a moderate (CV% < 50%) to slightly high (CV% > 50%) inter-animal variability in plasma levels. The inter-variability in PK parameters was also acceptable as it was moderate whatever the tested dose level. Additionally, as already observed in the previous cattle study in laboratory conditions with mdc-STM-001 tested at the lowest dose of 0.6 mg ivermectin/kg [34], it is worth noting that the initial plasma concentration (*i.e.,* burst or peak level) burst remained under control range even at the high dose of 1.5 mg/kg (Cmax mean [range]: 81.7 ng/mL [25–121]). This outcome is reassuring from a safety perspective. Indeed, the observed Cmax remains far below the mean safety threshold of 248 ng/mL previously reported in healthy volunteers administered escalating oral doses of ivermectin up to 120 mg (approximately 1404-2000 μg/kg) [23]. On top of the adequate safety profile, the ivermectin release performance and the good bio-resorption of the mdc-STM-001 depot overtime, the positive outcome is that a dose proportionality was demonstrated between 1.0 and 1.5 mg/kg for Cmax and AUC. In this context, efficacy depends not only on peak concentration (Cmax or burst) or total exposure (AUC) but also on how long drug levels remain above the thresholds required to kill mosquitoes. However, a consistent increase in effect duration was not shown at the mdc-STM-001 high dose (1.5 mg/kg) across species. While a longer effect is observed for *An. coustani* and *An. squamosus,* no meaningful change is observed for *An. funestus* and a similar duration is suggested for *An. rufipes.* This indicates that the dose-response relationship is not uniform and likely reflects species-specific susceptibility and non-linear pharmacokinetic-pharmacodynamic relationships. To evaluate the proportionality between 0.6 and 1.5 mg/

### Abundance and diversity of Anopheles

The species composition of collected mosquitoes reflected the ecological diversity of vectors in the area. In western Africa, and particularly in Burkina Faso, *Anopheles gambiae s.s.*, *An. coluzzii*, *An. arabiensis*, and *An. funestus* are recognized as the primary malaria vectors [53–55]. However, recent entomological surveys have revealed heterogeneity in mosquito community composition across ecological settings, with secondary vector species contributing to the overall diversity and abundance. Some studies reported that although *An. gambiae* complex members and *An. funestus* remain dominant, other species such as *An. coustani, An. rufipes, An. squamosus, An. nili, An. pharoensis* can represent an important fraction of collected mosquito [27, 56]. The secondary vector *Anopheles* coustani [57] was the most abundant in our collections. Traditionally considered of minor epidemiological relevance because of its marked zoophilic tendencies, emerging evidence suggests that *An. coustani* may occasionally display unexpected levels of anthropophagy. Human blood feeding, *Plasmodium* infections, and involvement in malaria transmission have now been reported in several African countries including Zambia, Kenya, Madagascar and Cameroun [58–63]. The predominance of *An. coustani* in our collections is particularly noteworthy and is consistent with increasing evidence that this species could play an important role in residual malaria transmission [56]. As the effectiveness of indoor interventions has increased, malaria transmission is increasingly maintained by vector species that bite outdoors, feed opportunistically on both humans and animals and avoid contact with insecticide-treated surfaces [64]. Such ecological traits allow secondary vectors to persist despite sustained control pressure and potentially compensate for declines in primary vectors. A similar perspective applies to *Anopheles rufipes* [56, 65] and *Anopheles squamosus* [66, 67] which were also abundant and among the most frequently collected in tent traps. Their abundance and increasing implication in transmission suggest that secondary vectors should no longer be overlooked. Consequently, surveillance programs and vector control strategies should increasingly consider the contribution of secondary vectors alongside the traditional focus on primary vectors.

Interestingly, when analyses focused on species diversity, treatment effects remained globally non-significant. The Shannon index further supports that species diversity remained stable across treatment groups and sampling rounds. This indicates that ivermectin exposure did not induce generalized behavioral deterrent effect at mosquito populations level. Overall, the results suggest that ivermectin’s primary impact is unlikely to operate through reduced or increased mosquito attraction. Likewise, the absence of treatment-related changes in species diversity suggests that ivermectin does not modify the composition of mosquito communities. Rather than favoring or disadvantaging particular vector species, ivermectin appears to exert its mosquitocidal activity after blood feeding without promoting shifts in species dominance or replacement within the mosquito populations.

### Long-lasting mosquitocidal efficacy of the ivermectin slow-release formulation

The present study demonstrates that the long-acting ivermectin formulation mdc-STM-001 induces potent and sustained mosquitocidal effects against wild *Anopheles* mosquitoes. By combining experimental rounds and integrating survival probability, cumulative mortality, median survival, hazard ratios, and dose-response data, our results provide evidence that the 1.0 and 1.5 mg ivermectin/kg subcutaneous doses reduce mosquito survival. The mosquitocidal effect was maintained for more than one month across the overall *Anopheles* population and persisted for more than two months in some species like *An. funestus* and *An. rufipes*. Both ivermectin doses increased early mosquito mortality compared with untreated, consistent with the expected pharmacokinetic profile in which the highest plasma concentrations occur shortly after administration, followed by gradual decrease of the rate of absorption from the subcutaneous depot [68]. The persistence of mortality beyond the initial peak exposure indicates that the mdc-STM-001 successfully maintains ivermectin concentrations above lethal thresholds for prolonged periods. Such long-lasting activity is rare with conventional ivermectin, whose efficacy usually declines few days after administration in cattle [69, 70]. Hazard ratios (HRs) provide a more formal quantification of effect size, demonstrating a consistent increase in mortality risk among mosquitoes exposed to treated cattle compared with controls. The observed effect sizes exceeded the threshold of 4recommended by WHO for evaluating endectocidal tools [52], supporting the potential of ivermectin long-acting formulation to maintain biologically relevant mosquito-killing activity over sustained periods. These findings are consistent with pharmacokinetic data for long-acting formulations, which indicate that lethal plasma concentrations can be maintained for approximately two months in cattle [34] . Dose-response analyses revealed marked interspecific differences in susceptibility to ivermectin, underscoring that the operational performance of long-acting formulations depends not only on systemic drug exposure but also on species-specific pharmacodynamic responses. For most suceptible species such as *An. funestus* and *An. rufipes*, the exposure achieved with the lower dose appeared sufficient to maintain effective mosquitocidal activity over a sustained period, suggesting limited added benefit from dose escalation once pharmacodynamic thresholds are reached. In contrast, less susceptible species *An. coustani* and *An. squamosus*, showed a greater benefit from the higher dose, indicating that increased systemic exposure may be required to achieve and maintain effective concentrations against species with lower ivermectin sensitivity. Therefore, the optimal dosing strategy for long-acting ivermectin formulations may depend on the composition of local vector populations and the relative susceptibility profiles of target species. The observed differences in susceptibility are consistent with previous studies reporting marked interspecific variation in ivermectin responsiveness among malaria vectors [71, 72]. While the underling mechanisms were not investigated here, differences in physiological or metabolic traits, as well as other species-specific biological characteristics, may contribute to the heterogeneous responses observed. Nevertheless, the dose-dependent differences should be interpreted with caution. Because each dose was evaluated using independent collections of wild mosquitoes sampled longitudinally, individuals exposed to similar ivermectin concentrations were not necessarily drawn from the same underlying populations. Seasonal changes in mosquito population structure, including variation in age, body size, physiological or metabolic status may all influence ivermectin susceptibility. It is also plausible that the 1.5 mg/kg high dose modified the relative abundance or persistence of bioactive ivermectin metabolites, thereby inducing greater mosquitocidal activity.

Despite some variability in pharmacokinetic profiles, both doses of the slow-release formulation were able to maintain biologically relevant ivermectin concentrations for extended periods across a broad range of *Anopheles* species. However, the observed outcomes suggest that dose escalation does not translate into uniform gains across species but rather interacts with species-specific pharmacodynamic sensitivity and may also be influenced by the age structure of field mosquito populations.

Overall, these results emphasize that the effect of ivermectin slow-release formulation is species-dependent and point out the importance of integrating species-specific pharmacodynamic, mosquito population structure and inter-individual pharmacokinetic variability in predicting ivermectin’s operational impact.

## Conclusion

Our findings revealed the potential of long-acting IVM formulation as a complementary malaria control tool, particularly by targeting outdoor and exophilic mosquito populations that are less affected by conventional interventions such as ITNs and IRS. The sustained-release profile of such formulation could reinforce long lasting mosquito control effect without requiring repeating the drug administration. Such a strategy represents one of the few currently explored approaches capable of directly targeting mosquitoes involved in residual outdoor transmission. Further field trials are requested to validate the impact of dosing strategies in malaria transmission and how the strategy can be integrated in the panel of vector control interventions implemented by national malaria control programs. Moreover, it is crucial to investigate the possible emergence of vector resistance to IVM, as this could compromise the efficacy of such promising formulation. Our findings provide basic insights on long-lasting IVM formulation highlighting its potential for epidemiological impact that is paving the way for a future phase-I trial in humans.

## Consent for publication

Not applicable

## Authors contribution

RKD, FAS and KM designed the study. COWO, ES, DDS and DSR collected the data. COWO, ES and DDS supervised field activities. COWO, SB, DDS and AP analyzed the data. COWO drafted the original manuscript. FAS, ES, ABS, MN, SLD, SB, EHAN, KM and RKD reviewed the manuscript. All authors read and approved the final manuscript.

## Data and code availability

Data and codes are available online via datasud link (ongoing)

