## Supplementary material for "Efficacy of a long lasting ivermectin formulation against malaria vectors in a semi-field trial in Bapla, Southwest of Burkina Faso": Supplementary files.docx

### Appendix

### Methods

#### Injection volumes of mdc-STM-001 per cattle

##### Supplementary file S1: Table 1. Injected volume (mL) per animal given the body weight (BW, kg) measured at day-2 before treatment

| Approach / Group | mdc-STM-001  Dose (mg/kg) | Animal ID | BW at arrival  (kg) | BW at day-2 injection (kg) | Injected volume (mL) |
| --- | --- | --- | --- | --- | --- |
| Hut / 1 | 1.0 | B4878 | 82.9 | 83.6 | 1.06 |
|  |  | B4880 | 92.1 | 82.8 | 1.05 |
|  |  | B4899 | 111.8 | 109.6 | 1.39 |
|  |  | B4881 | 112.2 | 119.8 | 1.52 |
|  |  | B4874 | 101.6 | 106.8 | 1.36 |
|  |  | B4863 | 103.2 | 111.6 | 1.42 |
| Hut / 2 | 1.5 | B4873 | 74.2 | 67.2 | 1.28 |
|  |  | B4866 | 90.6 | 85.4 | 1.62 |
|  |  | B4870 | 106.2 | 110 | 2.09 |
|  |  | B4892 | 117.2 | 116.6 | 2.22 |
|  |  | B4879 | 86.4 | 83.8 | 1.59 |
|  |  | B4872 | 91.6 | 102.4 | 1.95 |
| Tent / 4 | 1.0 | B4887 | 94.4 | 84.6 | 1.07 |
|  |  | B4883 | 100.2 | 101.4 | 1.29 |
|  |  | B4862 | 119.6 | 124.6 | 1.58 |
|  |  | B4898 | 119 | 130.2 | 1.65 |
|  |  | B4871 | 106.8 | 111.8 | 1.42 |
|  |  | B4864 | 108.4 | 107 | 1.36 |
| Tent / 5 | 1.5 | B4896 | 93.4 | 92.6 | 1.76 |
|  |  | B4886 | 97.6 | 106.6 | 2.03 |
|  |  | B4895 | 117 | 119.8 | 2.28 |
|  |  | B4885 | 120.6 | 114.2 | 2.17 |
|  |  | B4867 | 97.4 | 95.4 | 1.81 |
|  |  | B4882 | 101.2 | 102.2 | 1.94 |

#### Statistical analysis

##### Coproscopic analysis

The number of individuals for parasite type including *Strongle, Strongyloides, Moniezia, Toxocara, Fasciola, Paramphistom, Tenia, Ascaris, Trichurus* was quantified across the different treatment groups (Control, IVM two doses tested). An analysis of deviance table using Type II Wald chi-square tests was performed to assess the statistical significance of treatment effects.

### Results

#### Cattle follow up

##### Supplementary file S1: Table 2. Monthly weight in kg for cattle at each weighing date

| **Date** | **Cattle ID** | **Treatment** | **Weight (Kg)** |
| --- | --- | --- | --- |
| 11/09/2021 | B4865 | control | 115.2 |
| 11/09/2021 | B4866 | mdc-STM-1.5 | 85.4 |
| 11/09/2021 | B4868 | control | 101.2 |
| 11/09/2021 | B4873 | mdc-STM-1.5 | 67.2 |
| 11/09/2021 | B4877 | control | 102.4 |
| 11/09/2021 | B4878 | mdc-STM-1.0 | 83.6 |
| 11/09/2021 | B4880 | mdc-STM-1.0 | 82.8 |
| 11/09/2021 | B4883 | mdc-STM-1.0 | 101.4 |
| 11/09/2021 | B4886 | mdc-STM-1.5 | 106.6 |
| 11/09/2021 | B4887 | mdc-STM-1.0 | 84.6 |
| 11/09/2021 | B4889 | control | 101.6 |
| 11/09/2021 | B4896 | mdc-STM-1.5 | 92.6 |
| 18/09/2021 | B4862 | mdc-STM-1.0 | 124.6 |
| 18/09/2021 | B4870 | mdc-STM-1.5 | 110 |
| 18/09/2021 | B4875 | control | 105.6 |
| 18/09/2021 | B4876 | control | 90.8 |
| 18/09/2021 | B4881 | mdc-STM-1.0 | 119.8 |
| 18/09/2021 | B4885 | mdc-STM-1.5 | 114.2 |
| 18/09/2021 | B4890 | control | 108.8 |
| 18/09/2021 | B4892 | mdc-STM-1.5 | 116.6 |
| 18/09/2021 | B4895 | mdc-STM-1.5 | 119.8 |
| 18/09/2021 | B4898 | mdc-STM-1.0 | 130.2 |
| 18/09/2021 | B4899 | mdc-STM-1.0 | 109.6 |
| 18/09/2021 | B4900 | control | 108.2 |
| 26/09/2021 | B4863 | mdc-STM-1.0 | 111.6 |
| 26/09/2021 | B4864 | mdc-STM-1.0 | 107 |
| 26/09/2021 | B4867 | mdc-STM-1.5 | 95.4 |
| 26/09/2021 | B4871 | mdc-STM-1.0 | 111.8 |
| 26/09/2021 | B4872 | mdc-STM-1.5 | 102.4 |
| 26/09/2021 | B4874 | mdc-STM-1.0 | 106.8 |
| 26/09/2021 | B4879 | mdc-STM-1.5 | 83.8 |
| 26/09/2021 | B4882 | mdc-STM-1.5 | 102.2 |
| 26/09/2021 | B4884 | control | 110.6 |
| 26/09/2021 | B4888 | control | 139 |
| 26/09/2021 | B4893 | control | 126.6 |
| 26/09/2021 | B4894 | control | 109.2 |
| 14/10/2021 | B4865 | control | 123.4 |
| 14/10/2021 | B4866 | mdc-STM-1.5 | 95.2 |
| 14/10/2021 | B4868 | control | 112.4 |
| 14/10/2021 | B4873 | mdc-STM-1.5 | 73.2 |
| 14/10/2021 | B4877 | control | 110 |
| 14/10/2021 | B4878 | mdc-STM-1.0 | 86.8 |
| 14/10/2021 | B4880 | mdc-STM-1.0 | 90.4 |
| 14/10/2021 | B4883 | mdc-STM-1.0 | 107 |
| 14/10/2021 | B4886 | mdc-STM-1.5 | 112.8 |
| 14/10/2021 | B4887 | mdc-STM-1.0 | 91.6 |
| 14/10/2021 | B4889 | control | 113 |
| 14/10/2021 | B4896 | mdc-STM-1.5 | 109.6 |
| 14/10/2021 | B4862 | mdc-STM-1.0 | 125.6 |
| 14/10/2021 | B4870 | mdc-STM-1.5 | 112.2 |
| 14/10/2021 | B4875 | control | 108 |
| 14/10/2021 | B4876 | control | 91.6 |
| 14/10/2021 | B4881 | mdc-STM-1.0 | 121.6 |
| 14/10/2021 | B4885 | mdc-STM-1.5 | 116 |
| 14/10/2021 | B4890 | control | 111.6 |
| 14/10/2021 | B4892 | mdc-STM-1.5 | 117.2 |
| 14/10/2021 | B4895 | mdc-STM-1.5 | 122.4 |
| 14/10/2021 | B4898 | mdc-STM-1.0 | 131.4 |
| 14/10/2021 | B4899 | mdc-STM-1.0 | 113.2 |
| 14/10/2021 | B4900 | control | 111.2 |
| 14/10/2021 | B4863 | mdc-STM-1.0 | 112 |
| 14/10/2021 | B4864 | mdc-STM-1.0 | 108.6 |
| 14/10/2021 | B4867 | mdc-STM-1.5 | 96.6 |
| 14/10/2021 | B4871 | mdc-STM-1.0 | 114 |
| 14/10/2021 | B4872 | mdc-STM-1.5 | 103.8 |
| 14/10/2021 | B4874 | mdc-STM-1.0 | 108 |
| 14/10/2021 | B4879 | mdc-STM-1.5 | 86.2 |
| 14/10/2021 | B4882 | mdc-STM-1.5 | 103.8 |
| 14/10/2021 | B4884 | control | 112.2 |
| 14/10/2021 | B4888 | control | 140.8 |
| 14/10/2021 | B4893 | control | 127.8 |
| 14/10/2021 | B4894 | control | 110.6 |
| 23/11/2021 | B4865 | control | 121.8 |
| 23/11/2021 | B4866 | mdc-STM-1.5 | 98.4 |
| 23/11/2021 | B4868 | control | 110.8 |
| 23/11/2021 | B4873 | mdc-STM-1.5 | 78.6 |
| 23/11/2021 | B4877 | control | 112.6 |
| 23/11/2021 | B4878 | mdc-STM-1.0 | 93.8 |
| 23/11/2021 | B4880 | mdc-STM-1.0 | 102.4 |
| 23/11/2021 | B4883 | mdc-STM-1.0 | 115.8 |
| 23/11/2021 | B4886 | mdc-STM-1.5 | 121.8 |
| 23/11/2021 | B4887 | mdc-STM-1.0 | 106.8 |
| 23/11/2021 | B4889 | control | 118.4 |
| 23/11/2021 | B4896 | mdc-STM-1.5 | 114.6 |
| 23/11/2021 | B4862 | mdc-STM-1.0 | 128.8 |
| 23/11/2021 | B4870 | mdc-STM-1.5 | 110.6 |
| 23/11/2021 | B4875 | control | 110.2 |
| 23/11/2021 | B4876 | control | 97.4 |
| 23/11/2021 | B4881 | mdc-STM-1.0 | 130.4 |
| 23/11/2021 | B4885 | mdc-STM-1.5 | 126.2 |
| 23/11/2021 | B4890 | control | 115.8 |
| 23/11/2021 | B4892 | mdc-STM-1.5 | 129.8 |
| 23/11/2021 | B4895 | mdc-STM-1.5 | 130.6 |
| 23/11/2021 | B4898 | mdc-STM-1.0 | 145.2 |
| 23/11/2021 | B4899 | mdc-STM-1.0 | 117.6 |
| 23/11/2021 | B4900 | control | 121.8 |
| 23/11/2021 | B4863 | mdc-STM-1.0 | 109.4 |
| 23/11/2021 | B4864 | mdc-STM-1.0 | 113 |
| 23/11/2021 | B4867 | mdc-STM-1.5 | 99.2 |
| 23/11/2021 | B4871 | mdc-STM-1.0 | 112.8 |
| 23/11/2021 | B4872 | mdc-STM-1.5 | 106.4 |
| 23/11/2021 | B4874 | mdc-STM-1.0 | 110.6 |
| 23/11/2021 | B4879 | mdc-STM-1.5 | 92.4 |
| 23/11/2021 | B4882 | mdc-STM-1.5 | 105.6 |
| 23/11/2021 | B4884 | control | 113.8 |
| 23/11/2021 | B4888 | control | 143.6 |
| 23/11/2021 | B4893 | control | 115.8 |
| 23/11/2021 | B4894 | control | 108.8 |
| 22/12/2021 | B4865 | control | 123 |
| 22/12/2021 | B4866 | mdc-STM-1.5 | 101.8 |
| 22/12/2021 | B4868 | control | 111.4 |
| 22/12/2021 | B4873 | mdc-STM-1.5 | 84 |
| 22/12/2021 | B4877 | control | 105.8 |
| 22/12/2021 | B4878 | mdc-STM-1.0 | 102 |
| 22/12/2021 | B4880 | mdc-STM-1.0 | 116.6 |
| 22/12/2021 | B4883 | mdc-STM-1.0 | 131.8 |
| 22/12/2021 | B4886 | mdc-STM-1.5 | 140.8 |
| 22/12/2021 | B4887 | mdc-STM-1.0 | 108.2 |
| 22/12/2021 | B4889 | control | 117.8 |
| 22/12/2021 | B4896 | mdc-STM-1.5 | 120.8 |
| 22/12/2021 | B4862 | mdc-STM-1.0 | 127.6 |
| 22/12/2021 | B4870 | mdc-STM-1.5 | 114.2 |
| 22/12/2021 | B4875 | control | 113.4 |
| 22/12/2021 | B4876 | control | 104.8 |
| 22/12/2021 | B4881 | mdc-STM-1.0 | 149,8 |
| 22/12/2021 | B4885 | mdc-STM-1.5 | 138.6 |
| 22/12/2021 | B4890 | control | 107 |
| 22/12/2021 | B4892 | mdc-STM-1.5 | 141 |
| 22/12/2021 | B4895 | mdc-STM-1.5 | 140.2 |
| 22/12/2021 | B4898 | mdc-STM-1.0 | 152.2 |
| 22/12/2021 | B4899 | mdc-STM-1.0 | 125.8 |
| 22/12/2021 | B4900 | control | 131.8 |
| 22/12/2021 | B4863 | mdc-STM-1.0 | 117.6 |
| 22/12/2021 | B4864 | mdc-STM-1.0 | 119.2 |
| 22/12/2021 | B4867 | mdc-STM-1.5 | 105.8 |
| 22/12/2021 | B4871 | mdc-STM-1.0 | 120.2 |
| 22/12/2021 | B4872 | mdc-STM-1.5 | 109 |
| 22/12/2021 | B4874 | mdc-STM-1.0 | 114.6 |
| 22/12/2021 | B4879 | mdc-STM-1.5 | 89.2 |
| 22/12/2021 | B4882 | mdc-STM-1.5 | 110 |
| 22/12/2021 | B4884 | control | 103.4 |
| 22/12/2021 | B4888 | control | 148.8 |
| 22/12/2021 | B4893 | control | 99.2 |
| 22/12/2021 | B4894 | control | 113.6 |

#### Pharmacokinetics profile of ivermectin in the cattle plasma

##### Supplementary S1: Fig. 1A. Individual concentration-time profiles of ivermectin in cattle plasma after single subcutaneous administration of mdc-STM-001 per approach and per dose level (semi-log scale)

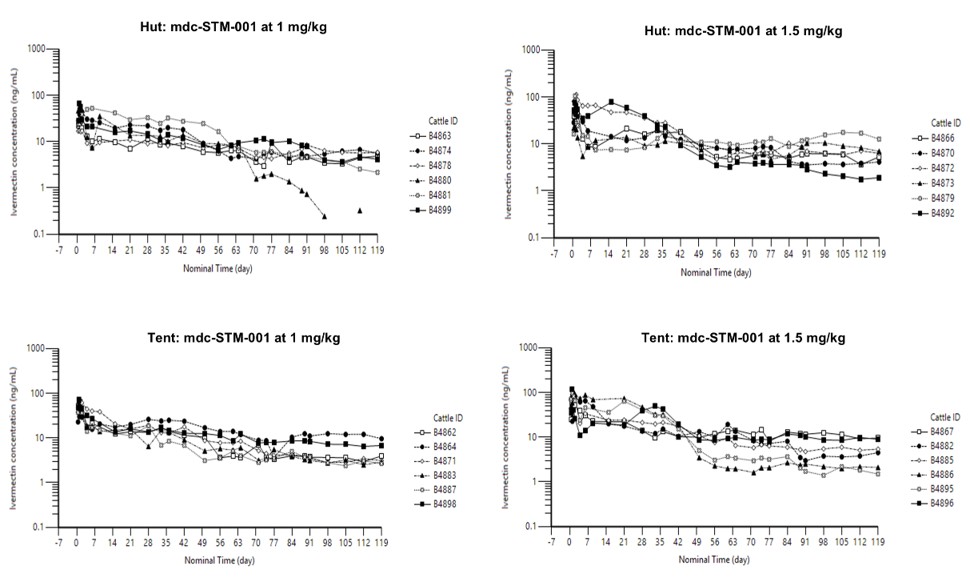

##### Supplementary file S1: Fig. 1B. Exposure approach effect: Mean (SD) concentration-time profiles of ivermectin in cattle plasma after single subcutaneous administration of mdc-STM-001 at 1.0 mg/kg

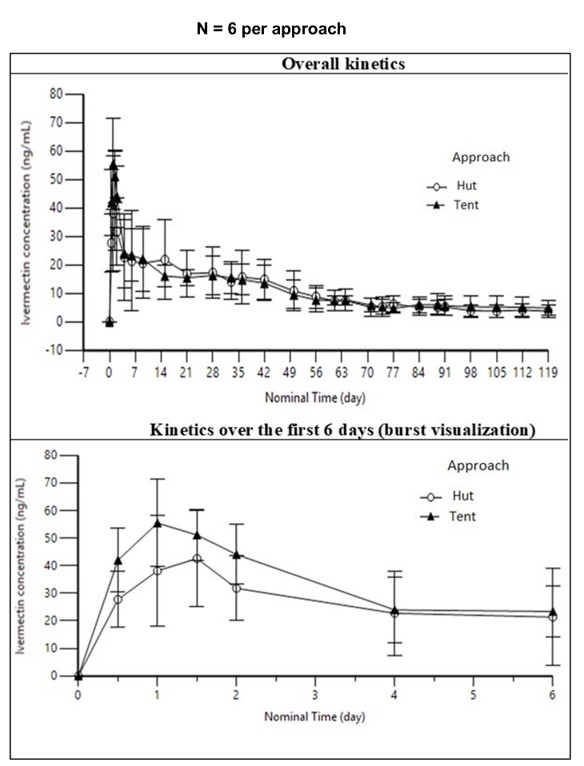

##### Supplementary file S1: Fig. 1C. Exposure approach effect: Mean (SD) concentration-time profiles of ivermectin in cattle plasma after single subcutaneous administration of mdc-STM-001 at 1.5 mg/kg

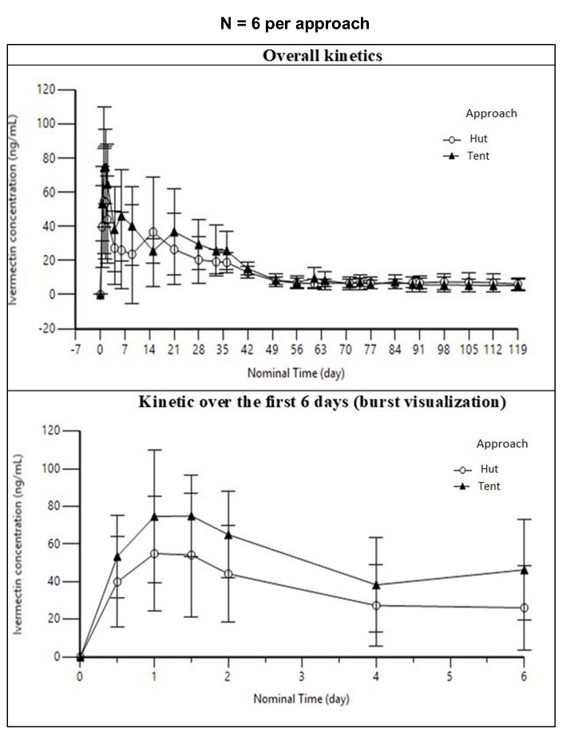

##### Supplementary file S1: Table 3. . Mean (SD) pharmacokinetic parameters of ivermectin in cattle plasma after single subcutaneous administration of mdc-STM-001 at 1.0 and 1.5 mg/kg; merged “hut” and “tent” data

**N=12 per dose level**

| Parameters | mdc-STM-001  1.0 mg/kg | mdc-STM-001  1.5 mg/kg |
| --- | --- | --- |
| Tmax* (h) | 24 (24 ; 48) | 36 (24 ; 360**) |
| Cmax (ng/mL) | 51.5 (17.1) | 81.7 (29.7) |
| Cmax/Dose (ng/mL/mg/kg) | 49.2 (16.6) | 53.9 (19.8) |
| AUC(0-28d) (h.ng/mL) | 14185 (5308) | 22557 (13115) |
| AUC(0-28d) /Dose  (h.ng/mL/mg/kg) | 13562 (5135) | 14813 (8494) |
| AUC(0-91d) (h.ng/mL) | 28156 (9442) | 38791 (13299) |
| AUC(0-91d) /Dose  (h.ng/mL/mg/kg) | 26957 (9302) | 25505 (8560) |
| AUC(0-119d) (h.ng/mL) | 31274 (10082) | 43040 (12013) |
| AUC(0-119d) /Dose  (h.ng/mL/mg/kg) | 29964 (10042) | 28310 (7701) |
| C28d (ng/mL) | 16.8 (7.51) | 25.0 (14.4) |
| C56d (ng/mL) | 8.25 (3.98) | 7.06 (3.25) |
| C91d (ng/mL) | 5.39 (2.93) | 6.34 (3.87) |
| C119d (ng/mL) | 4.71 (2.20) | 5.79 (3.39) |

SD: standard deviation.

*Median (min; max). Tmax: time to reach Cmax. Cmax : maximal plasma concentration. Cmax/Dose: dose-normalized Cmax. AUC(0-28d): area under curve from time 0 to 28 days. AUC(0-28d)/Dose: dose-normalized AUC(0-28d). AUC(0-91d): area under curve from time 0 to 91 days. AUC(0-91d)/Dose: dose-normalized AUC(0-91d). AUC(0-119d): area under curve from time 0 to 119 days (last common time-point for all animals). AUC(0-119d)/Dose: dose-normalized AUC(0-119d). C28d: plasma concentration 28 days post injection. C56d: plasma concentration 56 days post injection. C91d: plasma concentration 91 days post injection. C119d: plasma concentration 119 days post injection.

**one animal (“case”) with Tmax at 360h (*i.e.,* 15 days post injection): its concentration at 360h (77.4 ng/mL) is similar to its concentration at 24 hours post injection (76.0 ng/mL).

##### Supplementary file S1: Table 4. Mean (SD) pharmacokinetic parameters of ivermectin in cattle plasma after single subcutaneous administration of mdc-STM-001 at 1.0 and 1.5 mg/kg; each “hut” and “tent” data

**N=6 per approach and per dose level**

| Parameters | mdc-STM-001  1.0 mg/kg | | mdc-STM-001  1.5 mg/kg | |
| --- | --- | --- | --- | --- |
|  | Case / Gr. 1 | Tent / Gr. 4 | Case / Gr. 2 | Tent / Gr. 5 |
| Tmax* (h) | 30 (24; 48) | 24 (24; 48) | 36 (24; 360**) | 30 (24; 48) |
| Cmax (ng/mL) | 44.8 (19.3) | 58.2 (12.8) | 70.1 (34.7) | 93.3 (20.4) |
| Cmax/Dose (ng/mL/mg/kg) | 42.2 (17.9) | 56.3 (12.9) | 46.7 (23.6) | 61.2 (13.2) |
| AUC(0-28d) (h.ng/mL) | 14377 (7277) | 13993 (2982) | 18996 (14156) | 26118 (12148) |
| AUC(0-28d)/Dose  (h.ng/mL/mg/kg) | 13628 (7069) | 13497 (2834) | 12497 (9200) | 17129 (7823) |
| AUC(0-91d) (h.ng/mL) | 28558 (12115) | 27754 (6998) | 33840 (14973) | 43741 (10300) |
| AUC(0-91d)/Dose  (h.ng/mL/mg/kg) | 27077 (11865) | 26836 (7038) | 22340 (9703) | 28669 (6558) |
| AUC(0-119d) (h.ng/mL) | 31293 (11969) | 31255 (8965) | 38619 (13853) | 47462 (8870) |
| AUC(0-119d)/Dose  (h.ng/mL/mg/kg) | 29687 (11822) | 30240 (9049) | 25526 (9031) | 31093 (5506) |
| C28d (ng/mL) | 17.3 (8.86) | 16.3 (6.71) | 20.5 (13.8) | 29.4 (14.8) |
| C56d (ng/mL) | 8.82 (4.00) | 7.67 (4.25) | 6.87 (2.87) | 7.25 (3.86) |
| C91d (ng/mL) | 5.18 (2.76) | 5.59 (3.34) | 7.04 (3.70) | 5.64 (4.25) |
| C119d (ng/mL) | 3.78 (2.28) | 4.85 (2.80) | 6.22 (3.64) | 5.36 (3.39) |

SD: standard deviation. Gr: Group.

*Median (min; max). Tmax: time to reach Cmax. Cmax : maximal plasma concentration. Cmax/Dose: dose-normalized Cmax. AUC(0-28d): area under curve from time 0 to 28 days. AUC(0-28d)/Dose: dose-normalized AUC(0-28d). AUC(0-91d): area under curve from time 0 to 91 days. AUC(0-91d)/Dose: dose-normalized AUC(0-91d). AUC(0-119d): area under curve from time 0 to 119 days (last common time-point for all animals). AUC(0-119d)/Dose: dose-normalized AUC(0-119d). C28d: plasma concentration 28 days post injection. C56d: plasma concentration 56 days post injection. C91d: plasma concentration 91 days post injection. C119d: plasma concentration 119 days post injection.

**one animal (“case”) with Tmax at 360h (*i.e.,* 15 days post injection): its concentration at 360h (77.4 ng/mL) is similar to its concentration at 24 hours post injection (76.0 ng/mL).

#### Explant analysis at study end

##### Supplementary file S1: Table 5. Ivermectin extraction results in each depot collected at study end

| Approach / Group | mdc-  STM-001  Dose (mg/kg) | Animal ID | IVM injected dose (mg) | Time of depot collection  (DAI) | IVM recovered dose (mg) | | IVM recovered dose (%) | |
| --- | --- | --- | --- | --- | --- | --- | --- | --- |
|  |  |  |  |  | Indiv | Mean (SD) | Indiv | Mean (SD) |
| Case / 1 | 1.0 | B4878 | 88.9 | 150 | 27.7 | 6.7 (10.6) | 31.2 | 7.0 (12.0) |
|  |  | B4880 | 92.4 | 150 | 0.0 |  | 0.0 |  |
|  |  | B4899 | 116.9 | 143 | 3.4 |  | 2.9 |  |
|  |  | B4881 | 124.6 | 143 | 2.7 |  | 2.1 |  |
|  |  | B4874 | 108.9 | 135 | 6.3 |  | 5.8 |  |
|  |  | B4863 | 117.1 | 135 | 0.0 |  | 0.0 |  |
| Case / 2 | 1.5 | B4873 | 106.2 | 150 | 24.7 | 26.9 (22.2) | 23.2 | 19.8 (13.7) |
|  |  | B4866 | 128.9 | 150 | 13.0 |  | 10.1 |  |
|  |  | B4870 | 163.4 | 143 | 62.2 |  | 38.1 |  |
|  |  | B4892* | 179.2 | 143 | NR |  | NR |  |
|  |  | B4879 | 121.0 | 135 | 30.0 |  | 24.8 |  |
|  |  | B4872 | 155.9 | 135 | 4.4 |  | 2.8 |  |
| Tent / 4 | 1.0 | B4887 | 91.0 | 150 | 30.3 | 19.1 (15.5) | 33.3 | 18.1 (15.7) |
|  |  | B4883 | 105.1 | 150 | 41.5 |  | 39.5 |  |
|  |  | B4862 | 126.3 | 143 | 1.1 |  | 0.9 |  |
|  |  | B4898 | 134.2 | 143 | 10.1 |  | 7.5 |  |
|  |  | B4871 | 117.7 | 135 | 24.5 |  | 20.8 |  |
|  |  | B4864 | 108.4 | 135 | 7.3 |  | 6.7 |  |
| Tent / 5 | 1.5 | B4896 | 143.2 | 150 | 12.7 | 26.7 (26.7) | 8.8 | 17.3 (17.6) |
|  |  | B4886 | 168.1 | 150 | 5.6 |  | 3.3 |  |
|  |  | B4895 | 171.4 | 143 | 5.1 |  | 3.0 |  |
|  |  | B4885 | 177.1 | 143 | 20.7 |  | 11.7 |  |
|  |  | B4867 | 146.3 | 135 | 43.1 |  | 29.5 |  |
|  |  | B4882 | 155.0 | 135 | 73.2 |  | 47.2 |  |

IVM: ivermectin. Indiv: individual. SD: standard deviation. DAI: Day After Injection.

* Explant not found. NR: No Result.

#### Coproscopy

Coproscopic examinations revealed the presence of gastrointestinal nematodes belonging to the genera of *Strongyles* and *Strongyloides*, as well as cestodes of the genera *Moniezia* and *Taenia*, and trematodes from the Paramphistom group.

The analysis of parasite dynamics over time revealed differences in treatment efficacy across parasite types. For gastrointestinal nematodes, particularly *Strongyles* and *Strongyloides*, both doses induced a reduction in parasite loads shortly after administration. This effect was sustained throughout the observation period, with parasite counts in the mdc-STM-1.0 and mdc-STM-1.5 groups remaining consistently lower than in controls (χ^2^ = 10.43, df = 2, *p* = 0.005). In contrast, no significant reductions were recorded for *Moniezia*, *Taenia* and *Paramphistom* (*p* > 0.05), with parasite loads remaining comparable across treated and control groups.

#### Mosquito abundance and diversity

##### Supplementary file S1: Table 6. Results of Anova for final model of genus density in tent

**(a) Tent**

|  | **Chisq** | **Df** | **Pr(>Chisq)** |
| --- | --- | --- | --- |
| (Intercept) | 109.3329616 | 1 | 0.0000000 |
| Treatment | 0.8426585 | 2 | 0.6561740 |
| Round | 136.0205527 | 3 | 0.0000000 |
| Genus | 773.3763715 | 3 | 0.0000000 |
| Treatment : Round | 5.3977718 | 6 | 0.4938974 |
| Treatment : Genus | 2.7172025 | 6 | 0.8434112 |
| Round : Genus | 155.1484195 | 9 | 0.0000000 |
| Treatment : Round : Genus | 6.4665611 | 16 | 0.9822269 |

**(b) Hut**

|  | **Chisq** | **Df** | **Pr(>Chisq)** |
| --- | --- | --- | --- |
| (Intercept) | 21.9666901 | 1 | 0.0000028 |
| Treatment | 0.1166709 | 2 | 0.9433335 |
| Round | 25.4200662 | 3 | 0.0000126 |
| Genus | 49.1799405 | 3 | 0.0000000 |
| Treatment : Round | 3.4133818 | 6 | 0.7554559 |
| Treatment : Genus | 1.5384496 | 6 | 0.9568997 |
| Round : Genus | 41.8919230 | 8 | 0.0000014 |
| Treatment : Round : Genus | 6.4182479 | 14 | 0.9548240 |

##### Supplementary file S1: Fig. 2. Diversity and abundance of other mosquitoes (*Aedes, Culex, Mansonia*)

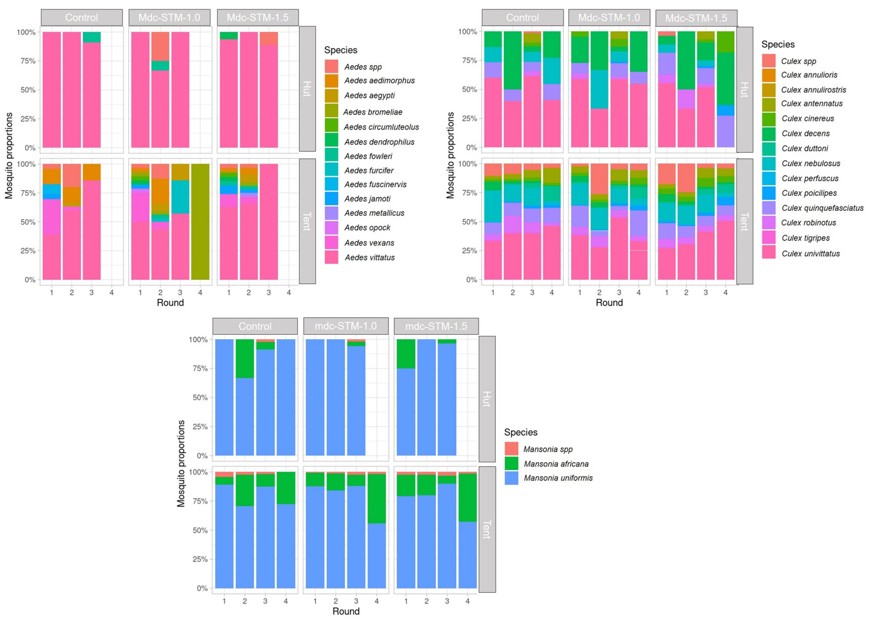

##### Supplementary file S1: Table 7. Contrast for *Anopheles* species density model in tent

| Contrast | Species | Round | Ratio | SE | LCL | UCL | z ratio | *p* value |
| --- | --- | --- | --- | --- | --- | --- | --- | --- |
| mdc-STM-1.0/Control | *An. spp* | 1 | 0.21 | 0.19 | 0.03 | 1.62 | -1.69 | 0.16 |
| mdc-STM-1.5/Control | *An. spp* | 1 | 0.19 | 0.13 | 0.04 | 0.85 | -2.46 | 0.02 |
| mdc-STM-1.0/Control | *An. coustani* | 1 | 1.43 | 0.24 | 0.98 | 2.09 | 2.15 | 0.05 |
| mdc-STM-1.5/Control | *An. coustani* | 1 | 1.80 | 0.30 | 1.24 | 2.61 | 3.54 | <0.001 |
| mdc-STM-1.0/Control | *An. flavicosta* | 1 | 1.94 | 1.74 | 0.26 | 14.24 | 0.74 | 0.67 |
| mdc-STM-1.5/Control | *An. flavicosta* | 1 | 1.16 | 1.05 | 0.16 | 8.62 | 0.17 | 0.96 |
| mdc-STM-1.0/Control | *An. funestus* | 1 | 3.14 | 0.81 | 1.77 | 5.58 | 4.42 | <0.001 |
| mdc-STM-1.5/Control | *An. funestus* | 1 | 2.19 | 0.58 | 1.22 | 3.94 | 2.98 | 0.005 |
| mdc-STM-1.0/Control | *An. gambiae* | 1 | 1.24 | 0.56 | 0.46 | 3.39 | 0.48 | 0.82 |
| mdc-STM-1.5/Control | *An. gambiae* | 1 | 1.27 | 0.59 | 0.45 | 3.57 | 0.52 | 0.81 |
| mdc-STM-1.0/Control | *An. nili* | 1 | 0.79 | 0.37 | 0.28 | 2.23 | -0.50 | 0.81 |
| mdc-STM-1.5/Control | *An. nili* | 1 | 1.23 | 0.63 | 0.4 | 3.84 | 0.41 | 0.86 |
| mdc-STM-1.0/Control | *An. pharoensis* | 1 | 1.73 | 0.58 | 0.82 | 3.65 | 1.63 | 0.18 |
| mdc-STM-1.5/Control | *An. pharoensis* | 1 | 1.60 | 0.56 | 0.73 | 3.49 | 1.33 | 0.31 |
| mdc-STM-1.0/Control | *An. pretoriensis* | 1 | 1.31 | 0.98 | 0.24 | 6.97 | 0.35 | 0.89 |
| mdc-STM-1.5/Control | *An. pretoriensis* | 1 | 0.76 | 0.57 | 0.14 | 4.00 | -0.36 | 0.89 |
| mdc-STM-1.0/Control | *An. rufipes* | 1 | 1.17 | 0.24 | 0.74 | 1.84 | 0.75 | 0.66 |
| mdc-STM-1.5/Control | *An. rufipes* | 1 | 1.14 | 0.24 | 0.71 | 1.82 | 0.63 | 0.74 |
| mdc-STM-1.0/Control | *An. squamosus* | 1 | 1.23 | 0.31 | 0.70 | 2.15 | 0.82 | 0.62 |
| mdc-STM-1.5/Control | *An. squamosus* | 1 | 2.48 | 0.59 | 1.45 | 4.22 | 3.78 | <0.001 |
| mdc-STM-1.0/Control | *An. welcomei* | 1 | - | - |  | - | - | - |
| mdc-STM-1.5/Control | *An. welcomei* | 1 | - | - |  | - | - | - |
| mdc-STM-1.0/Control | *An. spp* | 2 | 0.48 | 0.27 | 0.14 | 1.68 | -1.30 | 0.33 |
| mdc-STM-1.5/Control | *An. spp* | 2 | 0.39 | 0.17 | 0.14 | 1.04 | -2.13 | 0.06 |
| mdc-STM-1.0/Control | *An. coustani* | 2 | 1.13 | 0.19 | 0.78 | 1.63 | 0.73 | 0.68 |
| mdc-STM-1.5/Control | *An. coustani* | 2 | 1.30 | 0.21 | 0.90 | 1.87 | 1.61 | 0.19 |
| mdc-STM-1.0/Control | *An. flavicosta* | 2 | 1.08 | 0.62 | 0.30 | 3.84 | 0.1 | 0.98 |
| mdc-STM-1.5/Control | *An. flavicosta* | 2 | 1.22 | 0.66 | 0.37 | 4.04 | 0.7 | 0.89 |
| mdc-STM-1.0/Control | *An. funestus* | 2 | 1.47 | 0.47 | 0.73 | 2.98 | 1.23 | 0.37 |
| mdc-STM-1.5/Control | *An. funestus* | 2 | 1.36 | 0.42 | 0.68 | 2.72 | 0.98 | 0.51 |
| mdc-STM-1.0/Control | *An. gambiae* | 2 | 1.60 | 0.74 | 0.57 | 4.50 | 1.02 | 0.49 |
| mdc-STM-1.5/Control | *An. gambiae* | 2 | 1.24 | 0.58 | 0.43 | 3.53 | 0.46 | 0.84 |
| mdc-STM-1.0/Control | *An. nili* | 2 | 1.04 | 0.99 | 0.13 | 8.59 | 0.05 | 0.99 |
| mdc-STM-1.5/Control | *An. nili* | 2 | 2.00 | 1.93 | 0.23 | 17.16 | 0.72 | 0.68 |
| mdc-STM-1.0/Control | *An. pharoensis* | 2 | 1.29 | 0.62 | 0.45 | 3.74 | 0.54 | 0.79 |
| mdc-STM-1.5/Control | *An. pharoensis* | 2 | 1.68 | 0.82 | 0.57 | 4.99 | 1.06 | 0.46 |
| mdc-STM-1.0/Control | *An. pretoriensis* | 2 | - | - |  | - | - | - |
| mdc-STM-1.5/Control | *An. pretoriensis* | 2 | - | - |  | - | - | - |
| mdc-STM-1.0/Control | *An. rufipes* | 2 | 1.27 | 0.26 | 0.80 | 2.02 | 1.16 | 0.40 |
| mdc-STM-1.5/Control | *An. rufipes* | 2 | 1.23 | 0.25 | 0.78 | 1.933 | 1.02 | 0.49 |
| mdc-STM-1.0/Control | *An. squamosus* | 2 | 1.31 | 0.26 | 0.83 | 2.04 | 1.34 | 0.31 |
| mdc-STM-1.5/Control | *An. squamosus* | 2 | 1.46 | 0.29 | 0.95 | 2.26 | 1.95 | 0.09 |
| mdc-STM-1.0/Control | *An. welcomei* | 2 | 0.99 | 1.08 | 0.09 | 11.25 | -0.01 | 0.99 |
| mdc-STM-1.5/Control | *An. welcomei* | 2 | 1.19 | 0.94 | 0.21 | 6.84 | 0.22 | 0.95 |
| mdc-STM-1.0/Control | *An. spp* | 3 | 2.82 | 2.13 | 0.53 | 15.05 | 1.38 | 0.29 |
| mdc-STM-1.5/Control | *An. spp* | 3 | 1.50 | 1.41 | 0.19 | 12.01 | 0.43 | 0.85 |
| mdc-STM-1.0/Control | *An. coustani* | 3 | 0.90 | 0.16 | 0.60 | 1.35 | -0.56 | 0.78 |
| mdc-STM-1.5/Control | *An. coustani* | 33 | 0.77 | 0.13 | 0.52 | 1.55 | -1.42 | 0.27 |
| mdc-STM-1.0/Control | *An. flavicosta* | 3 | 1.38 | 0.64 | 0.50 | 3.85 | 0.70 | 0.69 |
| mdc-STM-1.5/Control | *An. flavicosta* | 3 | 1.35 | 0.064 | 0.47 | 3.89 | 0.64 | 0.73 |
| mdc-STM-1.0/Control | *An. funestus* | 3 | 0.78 | 0.22 | 0.42 | 1.45 | -0.88 | 0.57 |
| mdc-STM-1.5/Control | *An. funestus* | 3 | 1.29 | 0.35 | 0.70 | 2.37 | 0.93 | 0.55 |
| mdc-STM-1.0/Control | *An. gambiae* | 3 | 1.24 | 0.42 | 0.58 | 2.65 | 0.64 | 0.73 |
| mdc-STM-1.5/Control | *An. gambiae* | 3 | 1.26 | 0.42 | 0.60 | 2.64 | 0.70 | 0.69 |
| mdc-STM-1.0/Control | *An. nili* | 3 | 0.77 | 0.72 | 0.10 | 6.19 | -0.27 | 0.93 |
| mdc-STM-1.5/Control | *An. nili* | 3 | 0.75 | 0.60 | 0.13 | 4.43 | -0.35 | 0.89 |
| mdc-STM-1.0/Control | *An. pharoensis* | 3 | 1.33 | 0.92 | 0.28 | 6.23 | 0.40 | 0.87 |
| mdc-STM-1.5/Control | *An. pharoensis* | 3 | 1.22 | 0.83 | 0.27 | 5.50 | 0.29 | 0.92 |
| mdc-STM-1.0/Control | *An. pretoriensis* | 3 | - | - |  | - | - | - |
| mdc-STM-1.5/Control | *An. pretoriensis* | 3 | - | - |  | - | - | - |
| mdc-STM-1.0/Control | *An. rufipes* | 3 | 1.12 | 0.22 | 0.73 | 1.73 | 0.59 | 0.76 |
| mdc-STM-1.5/Control | *An. rufipes* | 3 | 0.98 | 0.19 | 0.63 | 1.51 | -0.10 | 0.98 |
| mdc-STM-1.0/Control | *An. squamosus* | 3 | 0.85 | 0.19 | 0.51 | 1.41 | -0.73 | 0.68 |
| mdc-STM-1.5/Control | *An. squamosus* | 3 | 1.18 | 0.27 | 0.71 | 1.97 | 0.73 | 0.68 |
| mdc-STM-1.0/Control | *An. welcomei* | 3 | 1.07 | 0.6 | 0.29 | 3.94 | 0.12 | 0.98 |
| mdc-STM-1.5/Control | *An. welcomei* | 3 | 0.94 | 0.61 | 0.22 | 4.02 | -0.09 | 0.98 |
| mdc-STM-1.0/Control | *An. spp* | 4 | 1.50 | 1.14 | 0.28 | 8.07 | 0.53 | 0.80 |
| mdc-STM-1.5/Control | *An. spp* | 4 | 1.49 | 1.14 | 0.27 | 8.15 | 0.52 | 0.81 |
| mdc-STM-1.0/Control | *An. coustani* | 4 | 1.35 | 0.27 | 0.86 | 2.11 | 1.47 | 0.25 |
| mdc-STM-1.5/Control | *An. coustani* | 4 | 1.31 | 0.27 | 0.83 | 2.07 | 1.34 | 0.30 |
| mdc-STM-1.0/Control | *An. flavicosta* | 4 | 0.93 | 0.38 | 0.37 | 2.32 | -0.17 | 0.96 |
| mdc-STM-1.5/Control | *An. flavicosta* | 4 | 0.96 | 0.42 | 0.37 | 2.52 | -0.09 | 0.98 |
| mdc-STM-1.0/Control | *An. funestus* | 4 | 0.99 | 0.34 | 0.46 | 2.15 | -0.02 | 0.99 |
| mdc-STM-1.5/Control | *An. funestus* | 4 | 0.89 | 0.29 | 0.43 | 1.85 | -0.35 | 0.89 |
| mdc-STM-1.0/Control | *An. gambiae* | 4 | 1.33 | 0.52 | 0.55 | 3.20 | 0.72 | 0.68 |
| mdc-STM-1.5/Control | *An. gambiae* | 4 | 0.96 | 0.40 | 0.38 | 2.42 | -0.10 | 0.98 |
| mdc-STM-1.0/Control | *An. nili* | 4 | - | - |  | - | - | - |
| mdc-STM-1.5/Control | *An. nili* | 4 | - | - |  | - | - | - |
| mdc-STM-1.0/Control | *An. pharoensis* | 4 | 0.84 | 1.42 | 0.03 | 23.29 | -0.10 | 0.91 |
| mdc-STM-1.5/Control | *An. pharoensis* | 4 | - | - |  | - | - | - |
| mdc-STM-1.0/Control | *An. pretoriensis* | 4 | - | - |  | - | - | - |
| mdc-STM-1.5/Control | *An. pretoriensis* | 4 | - | - |  | - | - | - |
| mdc-STM-1.0/Control | *An. rufipes* | 4 | 1.06 | 0.22 | 0.67 | 1.68 | 0.30 | 0.92 |
| mdc-STM-1.5/Control | *An. rufipes* | 4 | 1.03 | 0.21 | 0.65 | 1.64 | 0.14 | 0.98 |
| mdc-STM-1.0/Control | *An. squamosus* | 4 | 1.71 | 0.57 | 0.82 | 3.57 | 1.31 | 0.19 |
| mdc-STM-1.5/Control | *An. squamosus* | 4 | 1.28 | 0.45 | 0.59 | 2.79 | 0.72 | 0.68 |
| mdc-STM-1.0/Control | *An. welcomei* | 4 | 1.01 | 0.42 | 0.40 | 2.59 | 0.04 | 0.99 |
| mdc-STM-1.5/Control | *An. welcomei* | 4 | 1.23 | 0.51 | 0.49 | 3.08 | 0.51 | 0.81 |

##### Mosquitocidal activity evaluation

##### Supplementary file S1: Table 8. Dose LC50 ratios for *Anopheles* species

| Species | Treatment | Mortaliry % | Estimate | Std. Error | t-value | p-value |
| --- | --- | --- | --- | --- | --- | --- |
| All *Anopheles* | mdc-STM-1.0/ mdc-STM-1.5 | 50/50 | 1.506099 | 0.0644989 | 7.846625 | 0.0e+00 |
| *An. coustani* | mdc-STM-1.0/ mdc-STM-1.5 | 50/50 | 1.5506757 | 0.0680665 | 8.0902566 | 0.0000000 |
| *An. funestus* | mdc-STM-1.0/ mdc-STM-1.5 | 50/50 | 1.3573891 | 0.2859396 | 1.2498762 | 0.2113448 |
| *An. rufipes* | mdc-STM-1.0/ mdc-STM-1.5 | 50/50 | 0.9876222 | 0.1827764 | -0.0677207 | 0.9460079 |
| *An. squamosus* | mdc-STM-1.0/ mdc-STM-1.5 | 50/50 | 1.7568230 | 0.2139085 | 3.5380686 | 0.0004031 |

##### Supplementary file S1: Table 9. Anova for drc model selection

| Species | MoelDf | Loglik | Df | LR value | p-value | AIC | BIC |
| --- | --- | --- | --- | --- | --- | --- | --- |
| All *Anopheles* | With dose (df=6) | -830.8032 | NA | NA | NA | 1669.606 | 1683.052 |
|  | Without dose (df=3) | -874.5653 | 1 | 87.52421 | 0 | 1755.131 | 1765.143 |
| *An. coustani* | With dose (df=6) | -562.3102 | NA | NA | NA | 1132.6204 | 1145.8732 |
|  | Without dose (df=3) | -615.5929 | 1 | 106.565381 | 0.0000000 | 1237.1858 | 1247.1254 |
| *An. funestus* | With dose (df=6) | -119.9570 | NA | NA | NA | 247.9140 | 260.3135 |
|  | Without dose (df=3) | -118.1026 | 1 | 3.708829 | 0.0541254 | 242.2052 | 251.5048 |
| *An. rufipes* | With dose (df=6) | -275.3911 | NA | NA | NA | 558.7823 | 571.6853 |
|  | Without dose (df=3) | -274.3443 | 1 | 2.093689 | 0.1479085 | 554.6886 | 564.3658 |
| *An. squamosus* | With dose (df=6) | -214.0083 | NA | NA | NA | 436.0165 | 448.6757 |
|  | Without dose (df=3) | -224.3073 | 1 | 20.598085 | 0.0000057 | 454.6146 | 464.1090 |
